# A mechanism for the elimination of IgE plasma cells

**DOI:** 10.1101/2022.05.11.491510

**Authors:** Adam K. Wade-Vallance, Zhiyong Yang, Jeremy B. Libang, Marcus J. Robinson, David M. Tarlinton, Christopher D. C. Allen

**Affiliations:** Biomedical Sciences Graduate Program, University of California, San Francisco, CA 94143, USA; Cardiovascular Research Institute, University of California, San Francisco, CA 94143, USA; Sandler Asthma Basic Research Center, University of California, San Francisco, CA 94143, USA; Department of Anatomy, University of California, San Francisco, CA 94143, USA; Department of Immunology & Pathology, Monash University, Melbourne, VIC 3004 Australia; Department of Immunology Discovery, Genentech, Inc., South San Francisco, CA 94080, USA

## Abstract

The proper regulation of IgE production safeguards against allergic disease, highlighting the importance of mechanisms that restrict IgE plasma cell (PC) survival. IgE PCs have unusually high surface B cell receptor (BCR) expression, yet the functional consequences of ligating this receptor are unknown. Here, we found that BCR ligation induced BCR signaling in IgE PCs followed by their elimination. In cell culture, exposure of IgE PCs to cognate antigen or anti-BCR antibodies induced apoptosis. IgE PC depletion correlated with the affinity, avidity, amount, and duration of antigen exposure and required the BCR signalosome components Syk, BLNK, and PLCγ2. In mice with a PC-specific impairment of BCR signaling, the abundance of IgE PCs was selectively increased. Conversely, BCR ligation by injection of cognate antigen or anti-IgE depleted IgE PCs. These findings establish a mechanism for the elimination of IgE PCs through BCR ligation. This has important implications for allergen tolerance and immunotherapy as well as anti-IgE monoclonal antibody treatments.

## INTRODUCTION

The production of IgE specific for environmental antigens is a critical driver of allergic disease. Normally, the production of IgE is tightly controlled and the abundance of IgE in serum is orders of magnitude less than that of IgG. In the past decade, technical advancements in the detection of rare IgE-expressing cells have rapidly expanded our knowledge of the mechanisms that limit IgE production.

IgE-expressing (IgE) B cell responses are constrained by multiple key regulatory mechanisms. The initial class-switch recombination to IgE requires adequate IL-4 and is profoundly inhibited by IL-21, leading to the generation of limited numbers of IgE B cells (Burgis and Gessner, 2007; Muller et al., 2008; Robinson et al., 2017; Yang et al., 2020). Once generated, IgE B cells exhibit a propensity to undergo rapid differentiation into IgE antibody-secreting cells (plasmablasts or plasma cells [PCs], which we collectively refer to hereafter as PCs for simplicity) (Erazo et al., 2007; Yang et al., 2012). IgE B cells are rare within germinal centers (GCs) and their relative abundance there further diminishes over time (He et al., 2013; Yang et al., 2012). In GCs, B cells undergo iterative cycles of Ig mutation and selection, resulting in antibody refinement and affinity maturation, and ultimately differentiate into long-lived PCs or memory B cells. In contrast, most IgE PCs are short-lived and are generated independently from GCs, while IgE memory B cells have not been reliably detected (Allen, 2022; He et al., 2013; Jimenez-Saiz et al., 2019; Yang et al., 2012).

The skewed cell fate trajectories of IgE B cells are driven by antigen-independent signaling of the IgE B cell receptor (BCR). When B cells were cultured in the absence of cognate antigen, the expression of membrane IgE (mIgE) resulted in enhanced PC differentiation compared with the expression of other BCR isotypes (Haniuda et al., 2016; Yang et al., 2016). In addition, genetic deficiency or pharmacological inhibition of proteins involved in BCR signal transduction, such as Syk, Btk, BLNK, and CD19, resulted in a decrease in the frequency of IgE PCs in cell culture without antigen (Haniuda et al., 2016; Yang et al., 2016). A recent CRISPR screen in cell culture (Newman and Tolar, 2021) further highlighted the contribution of BCR signaling components to IgE PC differentiation without antigen. Overall, these data are consistent with a model that the IgE BCR has antigen-independent activity that promotes PC differentiation (Haniuda et al., 2016; Yang et al., 2016).

Based on this model, we predicted that mice with impaired BCR signal transduction would have reduced IgE PC differentiation, which would result in the enhanced differentiation and/or maintenance of IgE GC B cells. While these mice indeed had increased frequencies of IgE B cells in GCs, unexpectedly they also had increased numbers of IgE PCs (Haniuda et al., 2016; Yang et al., 2016). Moreover, a recent mutagenesis screen in mice found that IgE production was increased when BCR signaling was impaired (SoRelle et al., 2021). One possibility is that the increases in IgE PCs in the context of *in vivo* BCR signaling impairments were due to the enhanced IgE GC B cell responses. Indeed, in BLNK-deficient mice, a sustained increase in the abundance of IgE PCs was observed, and Ig sequencing four weeks after immunization showed somatic mutations consistent with a GC origin (Haniuda et al., 2016). However, we also observed an increase in IgE PCs early after immunization of mice with BLNK-deficiency or with heterozygous mutations in *Syk* (Yang et al., 2016), suggesting these PCs may be generated through the extrafollicular pathway, independently from GCs. We confirmed that most IgE PCs expressed unmutated, germline Ig sequences, consistent with a GC-independent origin, 9 days after immunization of mice with *Syk*-heterozygous activated B cells (Yang et al., 2016). The mechanism by which impairments of BCR signal transduction led to increases in both GC-dependent and GC-independent IgE PC responses *in vivo* remains unclear.

One possibility to account for these findings is that BCR signaling has direct effects on differentiated IgE PCs rather than only their IgE B cell precursors. While the differentiation of B cells into PCs is often thought to coincide with a shift from the expression of membrane Ig (mIg) to secreted Ig (sIg), we and others previously reported that IgE PCs upregulated mIg expression compared with IgE B cells (He et al., 2013; Ramadani et al., 2017; Yang et al., 2012). High mIg expression has also been observed on IgM and IgA PCs, and these BCRs were signaling-competent (Blanc et al., 2016; Pinto et al., 2013). Whether the BCR on IgE PCs is functional has not been directly examined. In addition, while previous studies focused on antigen-independent signaling of the IgE BCR (Haniuda et al., 2016; Yang et al., 2016), antigen stimulation of the BCR would be relevant to real-world immunizations, allergen exposures, allergen immunotherapies, and parasitic worm infections.

Here, we report that high surface mIg expression on IgE PCs corresponds with elevated phosphorylation of Syk, BLNK, and PLCγ2 after stimulation with antigen. Cell culture studies revealed that this BCR signaling depleted antigen-specific IgE PCs in a manner that corresponded to the strength and duration of BCR stimulation. Consistent with the induction of apoptosis in IgE PCs after cognate antigen exposure, BCR ligation led to caspase-3 activation in IgE PCs, followed by cell death, which was rescued by *Bcl2* overexpression. Antigen-induced IgE PC elimination was also blocked by inhibition of Syk or genetic deficiency in *Blnk* or *Plcg2*, highlighting the importance of the BCR signalosome to this process. *In vivo*, heterozygosity for *Syk* specifically in PCs resulted in an increase in the absolute number and relative frequency of IgE PCs. Conversely, ligating mIgE on IgE PCs with cognate antigen or a monoclonal antibody led to the robust depletion of IgE PCs. Based on these findings, we propose that IgE PCs are intrinsically regulated by BCR signaling-induced apoptosis.

## RESULTS

### IgE PCs have heightened BCR expression and signaling

Previous studies in mice revealed that IgE PCs, in addition to expressing large amounts of sIg, have markedly elevated surface mIg expression compared with IgE GC B cells and IgG1 PCs (He et al., 2013; Yang et al., 2012). However, the relative surface mIg abundance on IgE- expressing cells compared with IgM-expressing cells has not been reported. We therefore directly compared the surface BCR expression on IgE PCs to GC B cells and PCs of various isotypes *in vivo* (see Figure S1 for gating strategy). To control for possible effects of the Ig variable region on surface BCR expression, we utilized B cells from B1-8i (hereafter referred to as B1-8) mice (Sonoda et al., 1997). These B cells express a knock-in heavy chain VDJ specific for the hapten 4-hydroxy-3-nitrophenyl (NP) when paired with Igλ light chains. To compare mIg expression *in vivo*, we adoptively transferred naïve Igλ-enriched B1-8 B cells into *Cd19^-/-^* recipients, immunized them with NP conjugated to chicken gamma globulin (NP-CGG) in alum adjuvant, and then analyzed the surface BCR expression on responding B1-8 cells in the draining lymph nodes (dLNs) by flow cytometry (see Table S1 for reagents). For equivalent comparison of surface mIg expression, we stained the Ig light chain (LC) on cells with an antibody to Igλ.

We observed that IgE PCs had the highest surface mIg expression among all isotypes of GC B cells and PCs examined, which was more than an order of magnitude higher than IgG1 PCs and IgE GC B cells and was slightly higher (1.5- to 2-fold) than IgM PCs and naïve B cells (Figure 1A).

**Figure 1.**
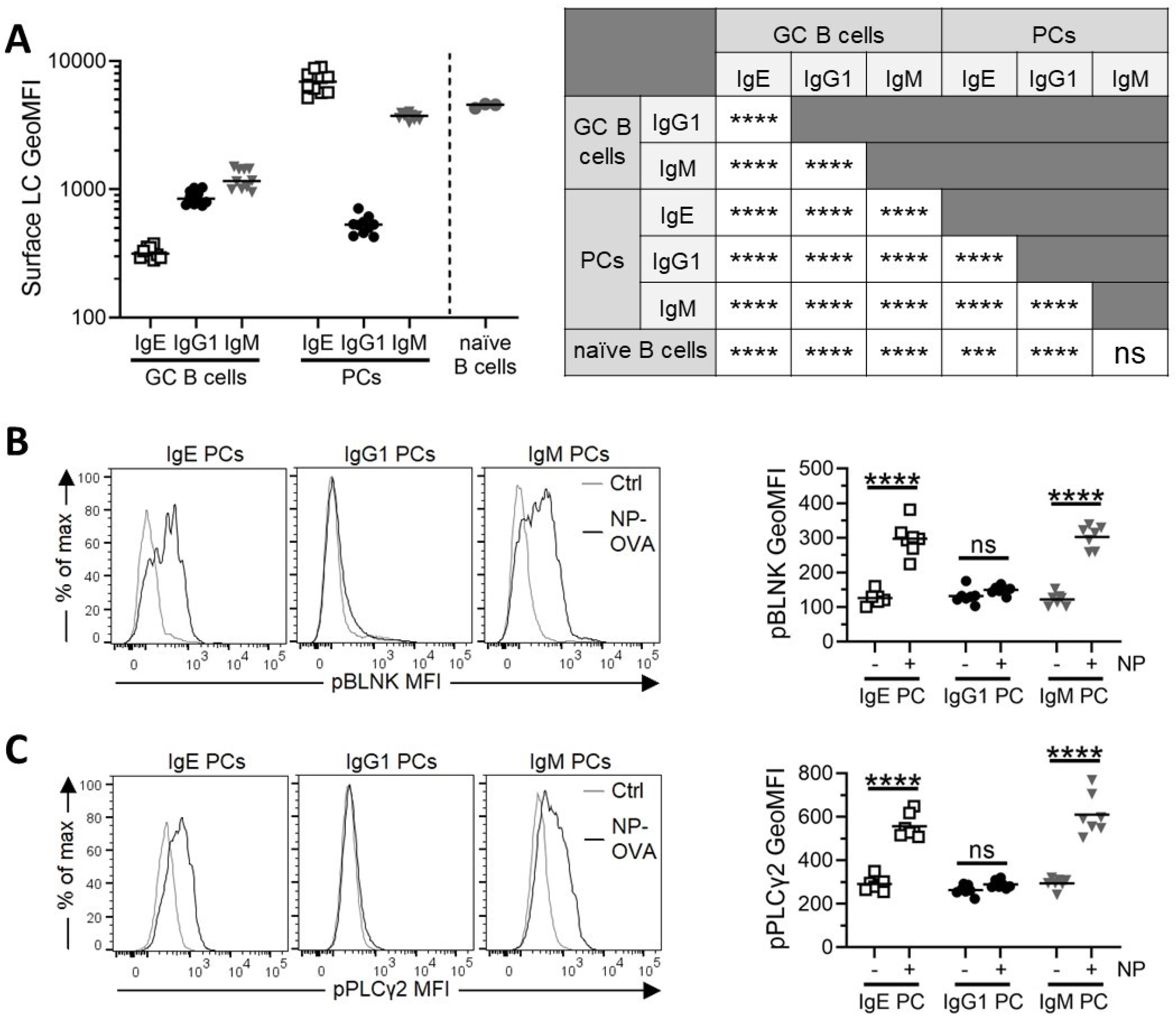
IgE PCs have high surface BCR expression and enhanced signaling in response to antigen. (A) Quantification of the surface expression of Igλ LC on the indicated cell types by flow cytometry. B1-8 B cells were adoptively transferred into *Cd19*^-/-^ mice, which were then immunized subcutaneously with NP-CGG in alum adjuvant. dLNs were collected 6 days later. After gating on congenic markers to identify transferred B1-8 cells, GC B cells were gated as B220^hi^CD138^−^PNA^+^CD38^−^IgD^−^ and PCs were gated as B220^int^CD138^+^ and negative for other isotypes, see Figure S1 for detailed gating strategy. Data for GC B cells and PCs are pooled from two independent experiments. For analysis of naïve B cells, splenocytes were isolated from separate B1-8 mice at the time of analysis and cells were gated as B220^hi^CD38^+^IgD^+^ Igλ^+^ and NP-binding. **(B-C)** Phosflow analysis of pBLNK **(B)** and pPLCγ2 **(C)** after *in vivo* BCR stimulation with representative histograms (left) and quantification (right). Mice were immunized subcutaneously with NP-CGG in alum adjuvant and then, 6.33-6.5 days later, were injected with NP- OVA 30 minutes prior to the collection of dLNs. PCs were gated as B220^int^CD138^+^ and negative for other isotypes. Data are pooled from three independent experiments. Dots represent samples from individual mice and bars represent the mean. n.s., not significant; **, P < 0.01; ***, P<0.001, ****, P < 0.0001 (one-way ANOVA with Tukey’s post-test comparing the mean of each group with the mean of every other group [A, table], one-way ANOVA with Sidak’s post-test comparing the means of pre-selected pairs of groups [B-C]).

We next tested whether ligation of mIg on IgE PCs would trigger an intracellular BCR signaling cascade as has been reported for IgM and IgA PCs (Blanc et al., 2016; Pinto et al., 2013). We injected antigen *in vivo* and examined the activation of intracellular signaling proteins by phosflow 30 minutes later. IgE and IgM PCs exhibited robust phosphorylation of BLNK (Figure 1B) and PLCγ2 (Figure 1C), whereas IgG1 PCs did not, indicating that IgE and IgM PCs can sense and respond to antigen *in vivo*.

To further explore BCR signaling in IgE PCs, we turned to an *in vitro* cell culture system. B cells were cultured with anti-CD40 (αCD40) and IL-4 to induce class-switch recombination and PC differentiation. For studies of BCR expression, we cultured Igλ-enriched B1-8 cells and detected surface BCR expression by flow cytometry with an antibody to the Igλ LC. In cell culture, we observed that IgE and IgM PCs had similar BCR expression, which was higher than IgG1 PCs (Figure S2A). Surface BCR expression on IgE PCs was also higher than on IgE B cells, which had several-fold lower surface BCR expression than IgG1 and IgM B cells (Figure S2A). Overall, these trends were similar to B1-8 cells differentiated in mice (Figure 1A), although we observed a further upregulation of surface BCR expression on IgE PCs generated *in vivo*.

To assess BCR signaling *in vitro*, we cultured polyclonal purified B cells, the vast majority of which express Igκ LC, and then ligated the BCRs with a polyclonal antibody to Igκ (αIgκ). Due to the increased numbers of IgE-expressing cells we were able to generate *in vitro*, we were able to assess the phosphorylation of Syk in addition to BLNK and PLCγ2. BCR ligation led to the phosphorylation of Syk (Figure S2B), BLNK (Figure S2C), and PLCγ2 (Figure S2D), which was more pronounced for IgE PCs and IgM PCs compared with IgG1 PCs, consistent with the relative amounts of surface BCR on these cells.

To determine whether BCR stimulation induced Ca^2+^ flux in IgE PCs, we took advantage of Verigem IgE reporter mice in which IgE-switched B cells and PCs express the Venus yellow fluorescent protein (Yang et al., 2012). Cultured Verigem B cells were loaded with the Ca^2+^- sensitive, ratiometric dye Indo-1 and were then stimulated with anti-BCR antibodies. Both IgE PCs and IgE B cells fluxed Ca^2+^, but IgE PCs exhibited stronger Ca^2+^ flux in response to BCR stimulation with either αIgE or αIgκ (Figure S2E). Together, these findings demonstrate that IgE PCs have elevated mIg which corresponds to increased signaling in response to BCR ligation.

### IgE PCs are preferentially lost in response to cognate antigen

We next sought to determine the functional consequence of BCR signaling in IgE PCs. In our Igλ-enriched B1-8 B cell culture system, approximately 50-80% of PCs expressed NP-specific BCRs. This allowed us to assess the responsiveness of NP-specific and non-specific IgE PCs in the same culture. When IgE PCs reached peak numbers at day 4.5, we added cognate antigen (NP conjugated to bovine serum albumin, NP-BSA) for BCR stimulation or control antigen (BSA). We then detected NP-specific cells by flow cytometry with NP-APC staining after fixation and permeabilization; this approach allowed us to detect antigen-specific PCs due to their abundant intracellular Ig, even if NP-BSA treatment might have blocked their surface mIg. After 24 hours of stimulation, we observed a profound decrease in the proportion of IgE PCs that were NP-binding, whereas there was no change in the proportions of IgG1 or IgM PCs that were NP-binding (Figure 2A). This result indicated that antigen recognition selectively disfavored antigen-specific IgE PCs. In an extensive timecourse analysis, the proportion of IgE PCs that were NP-binding decreased over time to a greater extent than for IgM PCs and IgG1 PCs (Figure 2B). Notably, the proportion of IgM and IgG1 PCs that were NP-specific transiently decreased at intermediate timepoints (6-12h), but by 24 hours had recovered. This phenomenon may have been due to antigen-induced BCR signaling in IgM and IgG1 B cells that promoted the differentiation of new IgM and IgG1 PCs, whereas IgE PC differentiation in cell culture is antigen-independent (Haniuda et al., 2016; Yang et al., 2016). To further assess if the observed reduction in the proportion of NP-binding IgE PCs after NP-BSA treatment was due to a decrease in the number of NP-binding cells or an increase in the number of non-antigen-binding cells, we enumerated these cells over the timecourse. The number of non-antigen specific IgE PCs was constant, whereas the number of NP-specific IgE PCs declined steadily with increasing duration of NP-BSA exposure (Figure 2C). These findings indicate that antigen-mediated stimulation of the BCR on IgE PCs leads to their loss over time.

**Figure 2.**
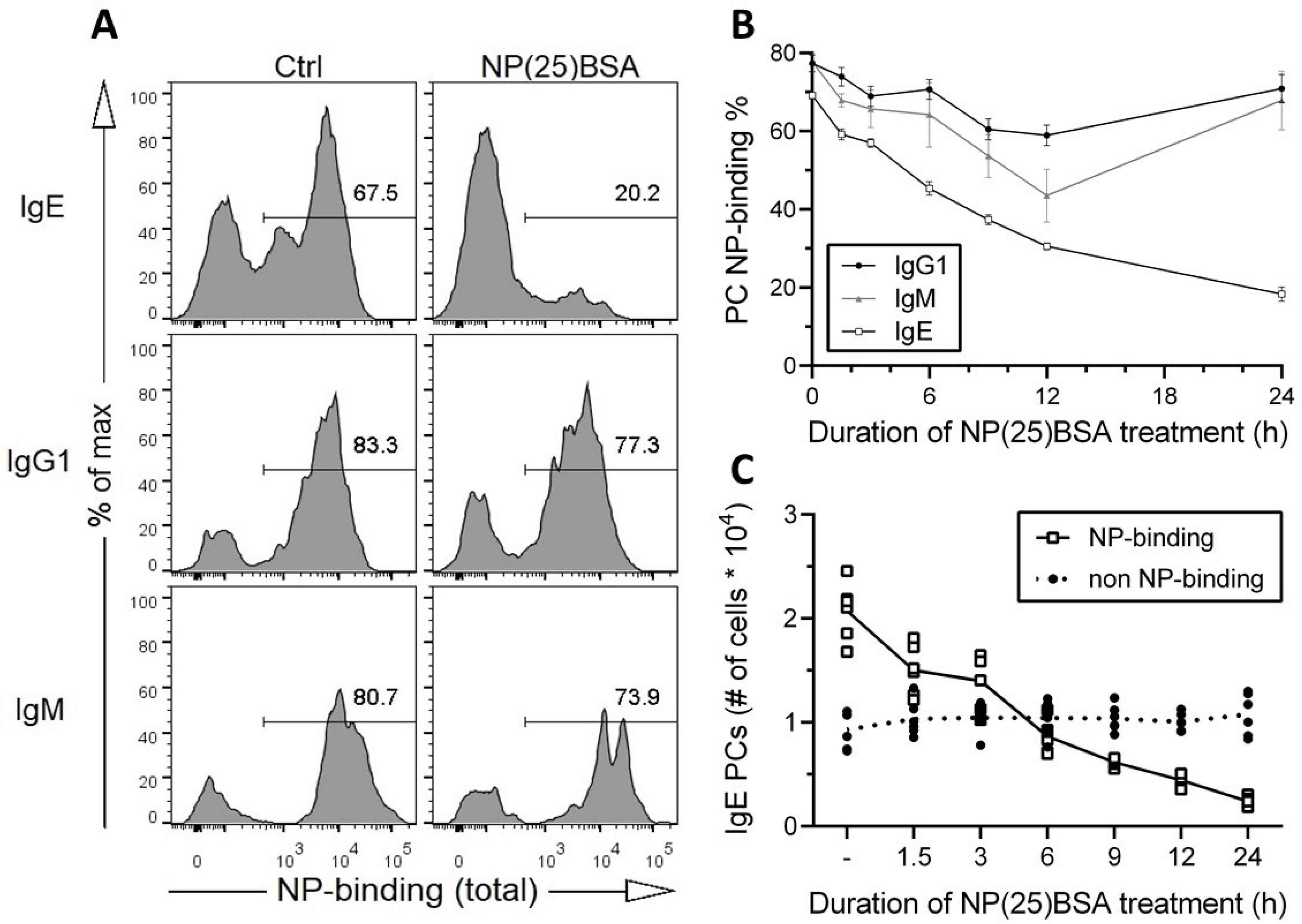
IgE PCs are preferentially depleted by cognate antigen over time. (A-C) Igλ-enriched B1-8 B cells were cultured for 5.5 days ± 6 hours. **(A)** Representative flow cytometry histograms showing the proportion of PCs (B220^int^CD138^+^) of the indicated isotypes with total (surface + intracellular) binding to NP-APC after 24 ± 3 hours of exposure to either NP(25)BSA (50ng/mL) or control antigen (BSA; 50ng/mL). **(B-C)** NP(25)BSA (50ng/mL) was added at the indicated timepoints, compared with a no antigen control (‘0h’ in [B] and ‘-’ in [C]), prior to analysis by flow cytometry, as in (A). **(B)** The proportion of PCs of each isotype that bound NP-APC is shown over time. Dots represent the mean of each group (cells from n=6 mice) and error bars reflect standard error. **(C)** The number of IgE PCs that were either NP-binding (white squares) or non-NP binding (black circles) is shown over time. Dots represent samples from individual mice. In (B-C), lines are drawn connecting the mean values. Results are representative of >10 independent experiments (A) or are pooled from two independent experiments (B, C).

### The loss of IgE PCs is proportional to the strength of BCR stimulation

We next investigated the relationship between the strength of BCR stimulation and the loss of IgE PCs in our B1-8 B cell culture system. BCR stimulation strength was modulated by altering the dose and valency of NP-BSA treatment for the final 24 hours of the culture and the proportion of IgE PCs that were NP-binding was determined by flow cytometry as above. The depletion of NP-specific IgE PCs was much more pronounced with high-valency NP(25)BSA than with low-valency NP(4)BSA and correlated with the amount of antigen added (Figure 3A). Since IgE PCs express higher surface BCR than IgG1 PCs, we reasoned that IgE PCs would be more sensitive to depletion following BCR stimulation. Therefore, we compared the NP-binding frequency of IgE, IgM, and IgG1 PCs following exposure to NP-BSA of varying doses and valency for 12 hours, a timepoint at which we had previously observed some reductions in PCs of all isotypes. Indeed, while all NP-specific PCs showed valency- and dose-dependent depletion, IgE PCs were depleted to a greater extent than IgG1 PCs, most notably with a high dose of low-valency NP(4)BSA or with a low dose of high-valency NP(25)BSA (Figure S3A). As IgM PCs were less abundant under these culture conditions, their responses showed greater variability, but compared with IgE PCs, they were similarly sensitive to low-valency NP(4)BSA, but less sensitive to high-valency NP(25)BSA. Therefore, IgE PCs exhibited enhanced elimination following BCR stimulation compared with IgG1 PCs and in some cases with IgM PCs.

**Figure 3.**
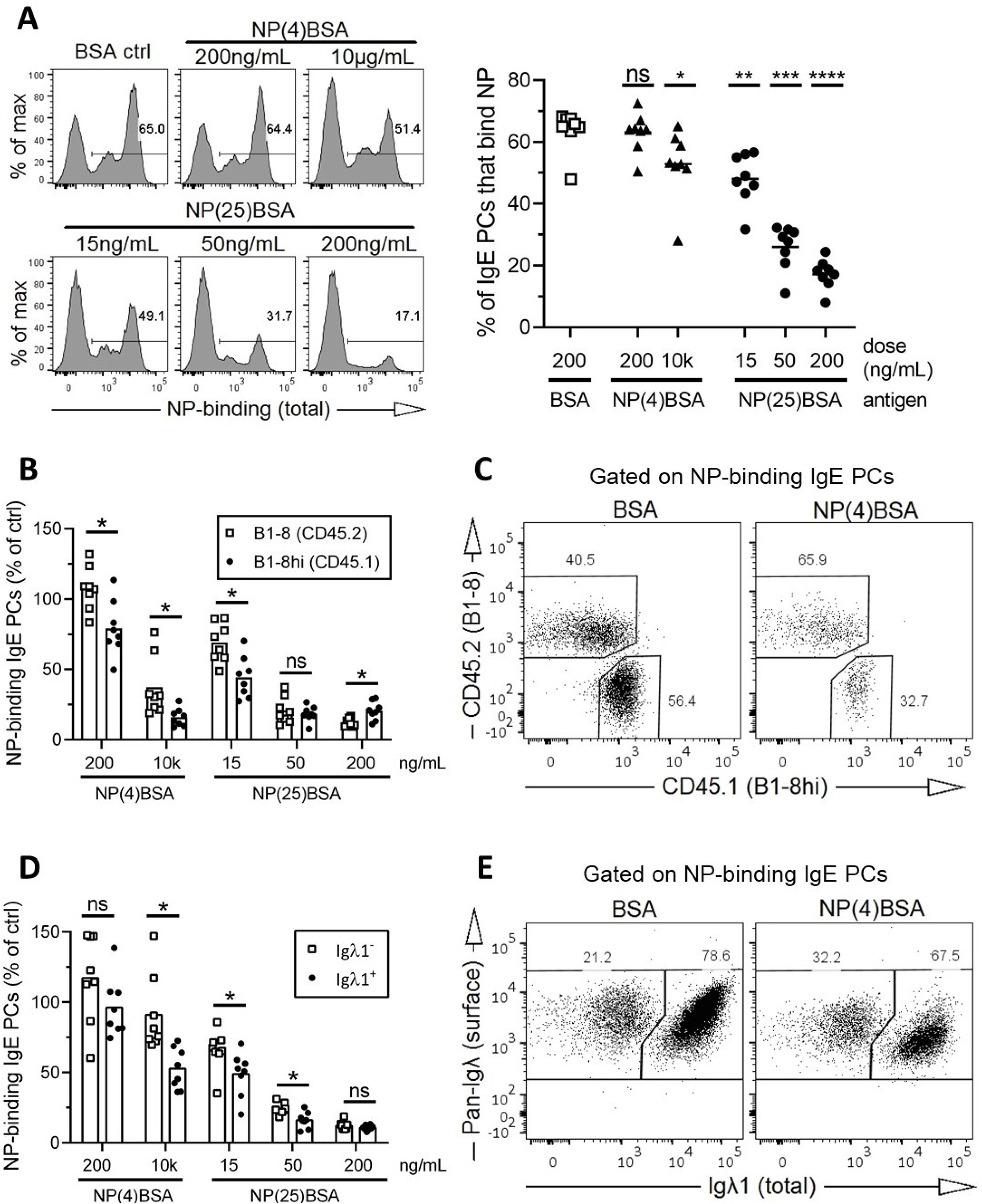
IgE PC depletion by cognate antigen is dose-dependent, affinity-dependent, and avidity-dependent. (A-E) Igλ-enriched B cells were cultured for 5.5 days ± 6 hours. The indicated antigens were added to the cultures 24 ± 3 hours prior to analysis by flow cytometry to determine the proportion of IgE PCs that bound to total (surface + intracellular) NP-APC. **(A)** Shown are representative histograms (left) and quantification (right) of the proportion of IgE PCs that were NP-binding after treatment with the indicated antigens. **(B)** Quantification of the number of NP-binding IgE PCs after incubation with the indicated antigens as a percentage of the number of NP-binding IgE PCs in control BSA-treated (200 ng/ml) wells. B cells were derived from B1-8 (white squares) and B1-8hi (black circles) mice and co-cultured, and then, as in (C), were identified by CD45 congenic markers. **(C)** Representative flow cytometry dot plots showing the proportion of NP-binding IgE PCs that were derived from B1-8 (CD45.2) or B1-8hi (CD45.1) donors after treatment with BSA as a control (200ng/mL) or with NP(4)BSA (10μg/mL). **(D)** Similar to B, except that only B1-8 cells were cultured and NP-binding IgE PCs were categorized as staining negatively (white squares) or positively (black circles) for intracellular Igλ1. **(E)** Representative flow cytometry dot plots showing the proportion of NP-binding IgE PCs that were Igλ_1_ ^−^or Igλ_1_ ^+^ after treatment with BSA or NP(4)BSA, as in (C). IgE PCs were gated as B220^int^CD138^+^IgE^+^. Dots in (A, B, D) show data points derived from cells from individual mice. n.s., not significant; *, P < 0.05; **, P < 0.01; ***, P < 0.001; ****, P < 0.0001 (one-way repeated measures ANOVA with Dunnett’s post-test comparing each group to the BSA ctrl [A] or unpaired [B] or paired [D] t tests with the Holm-Sidak correction for multiple comparisons). Bars represent the mean. Data are representative of (A, left panels; C; E) or pooled from (A, right panel; B; D) two independent experiments.

To determine how the affinity and avidity of antigen binding to the BCR affected cognate IgE PC depletion, we took advantage of a B1-8 line with a point mutation that confers 10-fold higher affinity to NP (B1-8hi) (Shih et al., 2002). We co-cultured Igλ-enriched cells from B1-8 and B1-8hi mice and then treated them with low- versus high-valency antigens at different doses as above for 24 hours. Low-valency or low-dose NP-BSA preferentially depleted B1-8hi IgE PCs (Figure 3B and C), indicating that the affinity of IgE PC mIg for antigen is most critical when antigens have low valency or are in low abundance.

As an additional affinity comparison, we took advantage of differences in Igλ LC expression among PCs in the B1-8 culture. A previous study identified two NP-binding populations of B1-8 cells that could be differentiated by their LC expression, one dominated by Igλ1 and the other by Igλ2 (Noviski et al., 2019). We found that PCs derived from Igλ-enriched B1-8 B cells were divisible into three populations with no NP-binding, low NP-binding, and high NP-binding (Figure S3B, left). Igλ1^+^ cells were predominantly high NP-binding with a smaller subset that was non-NP-binding, whereas the Igλ1^−^ (likely Igλ2^+^) cells were almost entirely intermediate NP-binding (Figure S3B, right). Among NP-binding PCs (gated as shown in Figure S3C, left), Igλ1^+^ staining clearly correlated with high NP-binding (Figure S3C, right), suggesting these cells would have increased sensitivity to cognate antigen. Indeed, low-valency or low-dose NP-BSA preferentially depleted Igλ1^+^ compared with Igλ1^−^ IgE PCs (Figure 3D and E), analogous to our results comparing higher-affinity B1-8hi to lower-affinity B1-8 IgE PCs. Taken together, these data provide evidence that the strength of BCR stimulation determines the extent of IgE PC depletion.

### IgE PCs undergo apoptosis following BCR stimulation in a cell-intrinsic manner

Our data above were consistent with a model in which stimulation of the BCR on IgE PCs resulted in signal transduction leading to apoptosis. However, these data did not exclude the possibility of indirect effects; for example, BCR stimulation could potentially affect the PC differentiation of precursor B cells. To formally establish whether BCR stimulation led to IgE PC apoptosis, we isolated pure populations of IgE PCs, non-IgE PCs, IgE B cells, and non-IgE B cells by sorting live *in vitro*-differentiated Verigem B cells by Venus fluorescence and CD138 staining (Figure S4A, left). Although IgG1- and IgM-expressing cells were indistinguishable at sorting, they were disentangled at the final analysis by staining with anti-isotype antibodies. To determine the effects of BCR stimulation, the sorted cell populations were cultured with αIgκ or control antibody for 24 hours. IgE PCs were depleted 4-fold by BCR stimulation, whereas IgG1 and IgM PCs showed a more modest 2-fold depletion (Figure 4A). This difference was not due to the expression of the Verigem reporter in IgE cells: in cultures of Verigem-heterozygous cells, the expression of mIgE from the Verigem allele did not promote, and in fact was slightly protective against, IgE PC elimination compared with the expression of mIgE from the wild-type allele (Figure S4B). We hypothesized that IgE PCs were undergoing apoptosis and sought to determine whether BCR stimulation led to increased caspase-3 activation. Indeed, we observed a 2-fold increase in active Caspase-3 (aCas3) staining among sorted IgE PCs treated with αIgκ (Figure 4B) but not among IgG1 or IgM PCs. Notably, BCR stimulation improved the survival of sorted IgE, IgG1, and IgM B cells and did not increase caspase-3 activation in these cells (Figure S4C and D), highlighting the different outcomes of BCR stimulation of IgE B cells versus IgE PCs. Taken together, these data provide evidence that BCR stimulation of IgE PCs results in caspase-3 activation and apoptosis.

**Figure 4.**
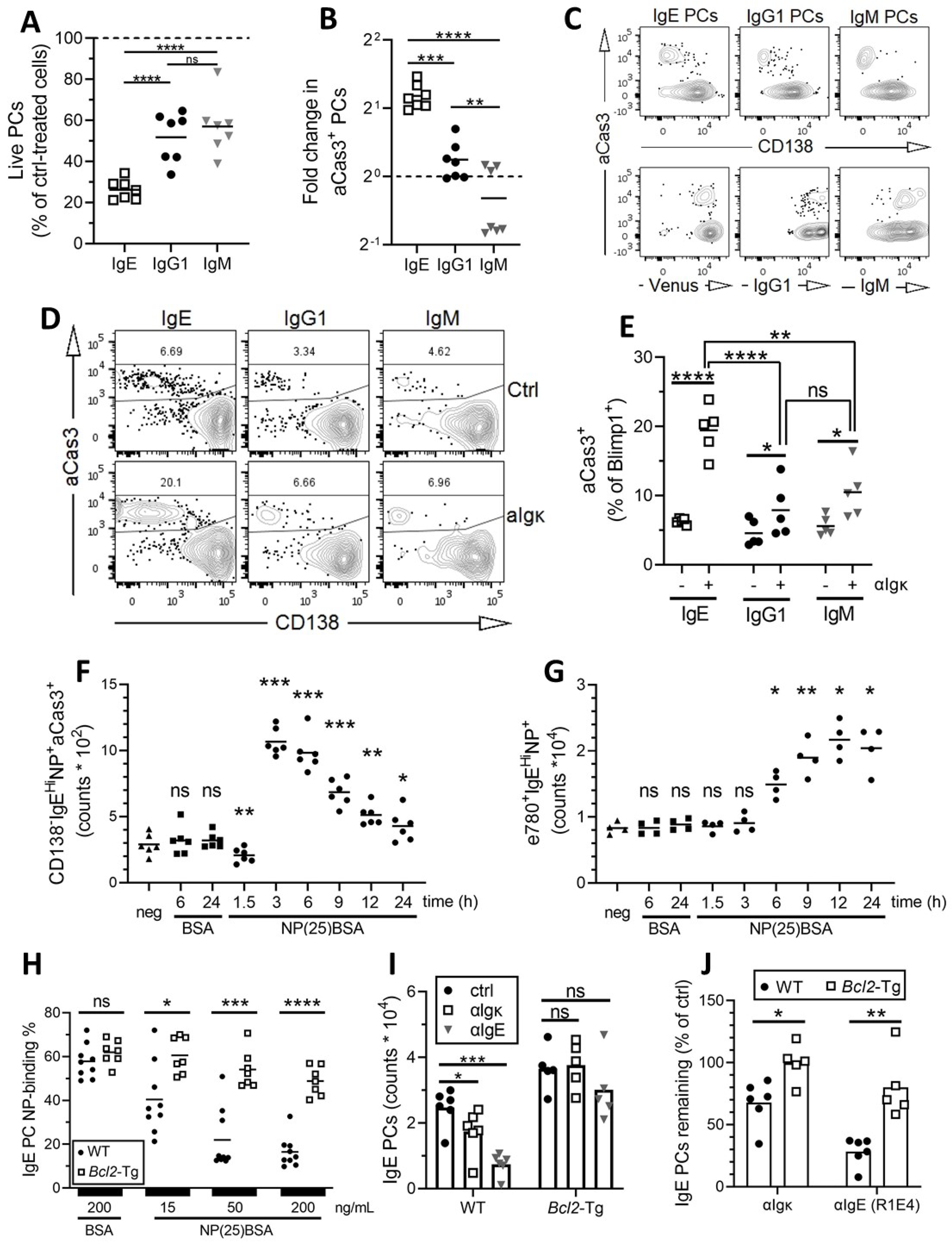
BCR stimulation results in IgE PC apoptosis. (A-C) Purified Verigem B cells were cultured for 4.5 days ± 6 hours, sorted by flow cytometry (see Figure S4A), returned to culture, treated with goat polyclonal IgG control or αIgκ antibody for 24 ± 3 hours, and then analyzed by flow cytometry. **(A)** Quantification of the number of live PCs after αIgκ treatment as a percentage of the number of live PCs after control IgG treatment for each isotype. The dotted line represents no change from control. **(B)** Quantification of the fold change in the proportion aCas3^+^ PCs after αIgκ treatment compared with control IgG treatment for each isotype. The dotted line represents no change from control. **(C)** Representative flow cytometry contour plots of aCas3 versus CD138 staining (top) or aCas3 versus fluorescence/isotype staining (bottom) for the indicated isotypes of PCs treated with αIgκ. **(D-E)** Purified B cells from Blimp1-YFP mice were cultured for 4.5 days and then treated with αIgκ (+) or goat IgG as a control (‘ctrl’ or ‘-’) for 6 hours prior to analysis by flow cytometry. **(D)** Representative flow cytometry contour plots of aCas3 versus CD138 staining for YFP^+^ cells of the indicated isotypes. **(E)** Quantification of the proportion of YFP^+^ cells of the indicated isotypes that were aCas3^+^. **(F-G)** Purified B cells from B1-8 mice were cultured for 5.5 days ± 6 hours. Cells were cultured with BSA (control) or NP(25)BSA at the indicated timepoints prior to analysis by flow cytometry, similar to Figure 2B and C. The number of CD138^−^IgE^hi^NP^+^aCas3^+^ cells **(F)** and e780^+^IgE^hi^NP^+^ cells **(G)** are shown. Cells were gated as shown in Figure S4E and F. **(H)** Quantification of the proportion of IgE PCs that were NP-binding after antigen treatment. Igλ-enriched B1-8 B cells from *Bcl2*-Tg (white squares) or control (WT, black circles) littermates were cultured for 5.5 days ± 6 hours and the indicated antigens were added for the final 24 ± 3 hours prior to analysis by flow cytometry. **(I-J)** Polyclonal B cells from *Bcl2-*Tg or WT littermates were purified and cultured for 5.5 days ± 6 hours. At D4.5 ± 6 hours, cells were treated with either αIgκ or αIgE for BCR stimulation, compared with control (goat IgG) treatment. Quantification of the absolute number of IgE PCs **(I)** or the relative percentage of IgE PCs remaining compared to control **(J)** after treatment as indicated. Dots represent data points derived from cells from individual mice. N.s., not significant; *, P < 0.05; **, P < 0.01; ***, P < 0.001; ****, P < 0.0001 (one-way ANOVA with Tukey’s post-test comparing each group to each other group [A, B], one-way matched ANOVA with the Sidak post-test comparing the indicated pairs of groups [E], one-way matched ANOVA with Dunnett’s post test comparing the mean of each group to the negative control [F, G], paired t tests with the Holm-Sidak correction for multiple comparisons [H, J], and two-way ANOVA matched within each genotype with the Sidak post test comparing the mean of each group to the control for each genotype [I]). IgE PCs were gated as B220^int^CD138^+^IgE^+^ (H-J). Bars represent the mean. Data are pooled from three (A-B), two (E-G), or four (H-J) independent experiments or are representative of three (C) or two (D) independent experiments.

Our studies of sorted PCs showed that apoptotic (aCas3^+^) PCs of all isotypes lost expression of CD138 (Figure 4C, top). These CD138^−^ aCas3^+^ cells were not contaminating B cells, because they retained similarly high intracellular antibody staining as live PCs (Figure 4C, bottom) and our post-sort purity routinely exceeded 99% (Figure S4A, right). Therefore, we identify CD138 downregulation as a general feature of PC apoptosis.

To further characterize PC apoptosis after BCR stimulation without cell sorting, we cultured cells from Blimp1-YFP-transgenic mice, in which YFP expression correlates with Blimp-1 expression and is therefore a marker of PC differentiation. The vast majority of YFP^+^ cells were also marked by CD138 (Figure 4D), indicating PC identity, but a small proportion of YFP^+^ cells was CD138^−^ and stained for aCas3, consistent with our prior observations that surface CD138 is lost during PC apoptosis. Treatment with αIgκ but not control antibody induced a large (3-4-fold) increase in the proportion of IgE^+^YFP^+^ cells that stained for aCas3 (Figure 4D and E), whereas only a slight increase in aCas3 staining was observed among IgG1^+^YFP^+^ cells and IgM^+^YFP^+^ cells. These results were consistent with our findings using sorted cells and provided further evidence that IgE PCs are uniquely sensitive to undergo apoptosis following BCR stimulation.

These observations prompted us to return to our timecourse experiments to examine the kinetics of BCR ligation-induced IgE PC apoptosis. We found that the addition of NP-BSA expanded a population of CD138^−^IgE^hi^ cells which was overwhelmingly (>95%) aCas3^+^ (Figure 4F and Figure S4E). This population spiked three hours after BCR stimulation and dwindled slowly thereafter. The proportion of these cells that were NP-binding was increased after NP- BSA treatment compared with control BSA treatment, consistent with the preferential induction of apoptosis in cognate antigen-specific IgE PCs (Figure S4E). Subsequent to the induction of aCas3, we observed an accumulation of CD138^−^ IgE^hi^ IgE PCs that stained with the fixable dye we used to exclude non-viable cells, suggesting these cells had completed apoptosis (Figure 4G). The frequency of NP-binding among these dead cells also increased following NP-BSA treatment (Figure S4F). Overall, these findings depict a progression of antigen-specific IgE PCs from live, to apoptotic, to dead following cognate antigen exposure.

Based on our previous finding that mice with an anti-apoptotic *Bcl2* transgene expressed in the B cell lineage had markedly increased numbers of IgE PCs (Yang et al., 2012), we sought to determine whether *Bcl2* overexpression could rescue IgE PCs from depletion after BCR stimulation. IgE PCs derived from *Bcl2-*Tg B1-8 B cells were strongly resistant to cognate antigen-induced elimination, at multiple doses of NP(25)BSA, relative to B1-8 IgE PCs derived from littermates lacking the *Bcl2* transgene (wildtype [WT]; Figure 4H). IgE PCs generated in cultures of polyclonal B cells from *Bcl2*-Tg mice without the B1-8 knock-in were also protected against elimination induced by αBCR antibodies relative to cells from WT littermates, showing no significant depletion even after exposure to αIgE, which depleted more than 75% of WT IgE PCs (Figure 4I and J). These data further support the model that BCR ligation induces apoptosis in IgE PCs, leading to their elimination.

### Antigen-induced IgE PC elimination requires the BCR signalosome

In Figure 1, we identified enhanced phosphorylation of BCR signalosome components following mIg stimulation of IgE PCs. Having observed that BCR stimulation led to IgE PC apoptosis, we next determined the contribution of BCR signalosome components to IgE PC elimination. First, we tested the role of Syk, an apical kinase in the BCR signaling phosphorylation cascade, and Btk, a kinase downstream of Syk, with the well-characterized pharmacological inhibitors PRT062607 and ibrutinib, respectively. Inhibition of either kinase dose-dependently rescued IgE PCs from BCR stimulation-induced elimination (Figure 5A and B). However, maximal Syk inhibition achieved a complete rescue, whereas maximal Btk inhibition had a partial effect. BLNK is an adaptor molecule critical for the activation of PLCγ2, which is important for Ca^2+^ flux and other downstream processes, and both BLNK and PLCγ2 were phosphorylated in IgE PCs following BCR stimulation (Figure 1). To test the contribution of these proteins to IgE PC depletion, we targeted the genes encoding these proteins by CRISPR- Cas9 ribonucleoprotein electroporation of B1-8 B cells. Targeting either *Blnk* or *Plcg2* completely blocked antigen-induced IgE PC elimination, whereas electroporation with Cas9 only (without a guide RNA) as a control did not (Figure 5 C and D). We observed similar results with multiple independent guides targeting either *Blnk* or *Plcg2* compared to a non-targeting sgRNA control (Figure S5A and B). These findings indicate that the mIg ligation-induced elimination of IgE PCs depends upon the activation of the BCR signalosome, with critical contributions from Syk, BLNK, and PLCγ2.

**Figure 5.**
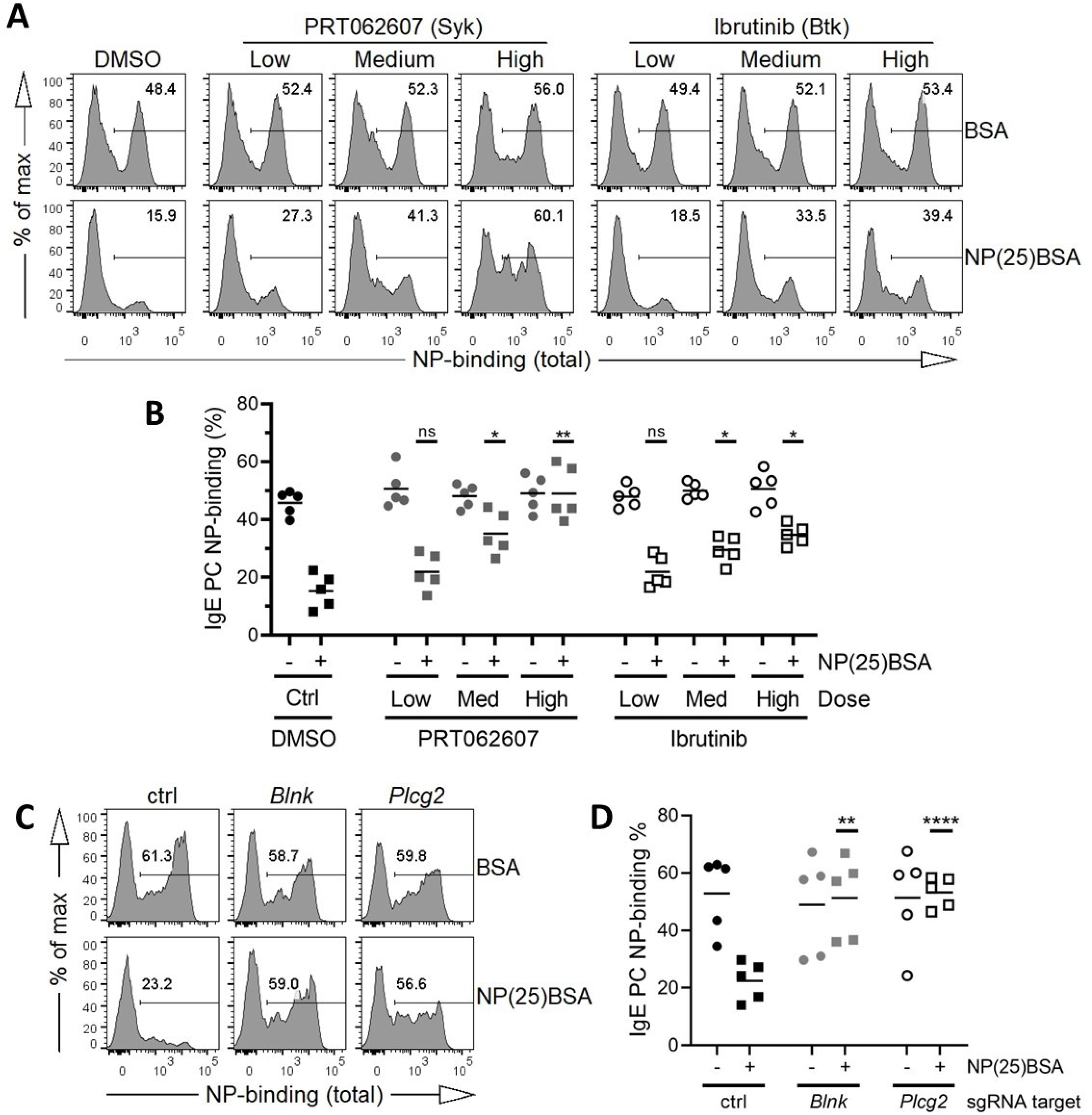
The BCR signalosome is required for mIg ligation-induced IgE PC elimination. (A-B) Igλ-enriched B1-8 B cells were cultured for 4.5 days ± 6 hours and then treated with NP(25)BSA or BSA (50 ng/mL) for 24 ± 3 hours prior to analysis. Shown are representative flow cytometry histograms **(A)** and quantification **(B)** of the proportion of IgE PCs that were NP-binding when cells were treated with the indicated inhibitors or vehicle control (DMSO) prior to incubation with antigen. See Table S2 for inhibitor concentrations. **(C-D)** Genes were targeted by electroporation with CRISPR-Cas9 ribonucleoproteins containing the indicated sgRNAs (or no sgRNA as a control) in cultured Igλ-enriched B1-8 B cells (see Methods for details). Representative flow cytometry histograms **(C)** and quantification **(D)** of the proportion of IgE PCs that were NP-binding after treatment with NP(25)BSA or BSA as a control (50-100ng/mL) for 24 ± 3 hours prior to analysis. IgE PCs were gated as B220^int^CD138^+^IgE^+^. Dots represent data points derived from cells from individual mice. n.s., not significant; *, P < 0.05; **, P < 0.01; ****, P < 0.0001 (one-way matched ANOVA with Dunnett’s post test comparing the mean of each NP(25)BSA-treated inhibitor condition to the NP(25)BSA-treated DMSO control [B] or each NP(25)BSA-treated sgRNA condition to the NP(25)BSA-treated no-sgRNA control [D]). Bars represent the mean. Data are representative of (A, C) or pooled from (B, D) two independent experiments.

### IgE PCs are constrained by BCR signaling *in vivo*

Having established that activation of the BCR signalosome led to the elimination of IgE PCs *in vitro*, we sought to confirm the importance of BCR signaling in restraining IgE PCs *in vivo*. To this end, we developed a genetic system where BCR signaling could be conditionally impaired exclusively in PCs. Specifically, we bred mice carrying a PC-restricted, tamoxifen-inducible Cre (BLIMP1-tdTomato-inducible Cre; BLTcre) (Robinson et al., 2022) and a single floxed *Syk* allele (*Syk*^flox/+^) (Saijo et al., 2003). To ensure Cre activity was restricted to the hematopoietic lineage and to produce matched group sizes sufficient for analysis, we generated bone marrow chimeras using BLTcre *Syk*^+/+^ donors as controls. Following immunization and tamoxifen treatment, IgE PCs were selectively increased in frequency (Figure 6A) and cell number (Figure 6B) in the dLNs of chimeras derived from *Syk*^flox/+^ donors compared with chimeras derived from *Syk*^+/+^ donors, whereas other isotypes of PCs and of GC B cells were unchanged (Figure 6B-C). These findings provide evidence for the selective regulation of IgE PCs *in vivo* by BCR signaling.

**Figure 6.**
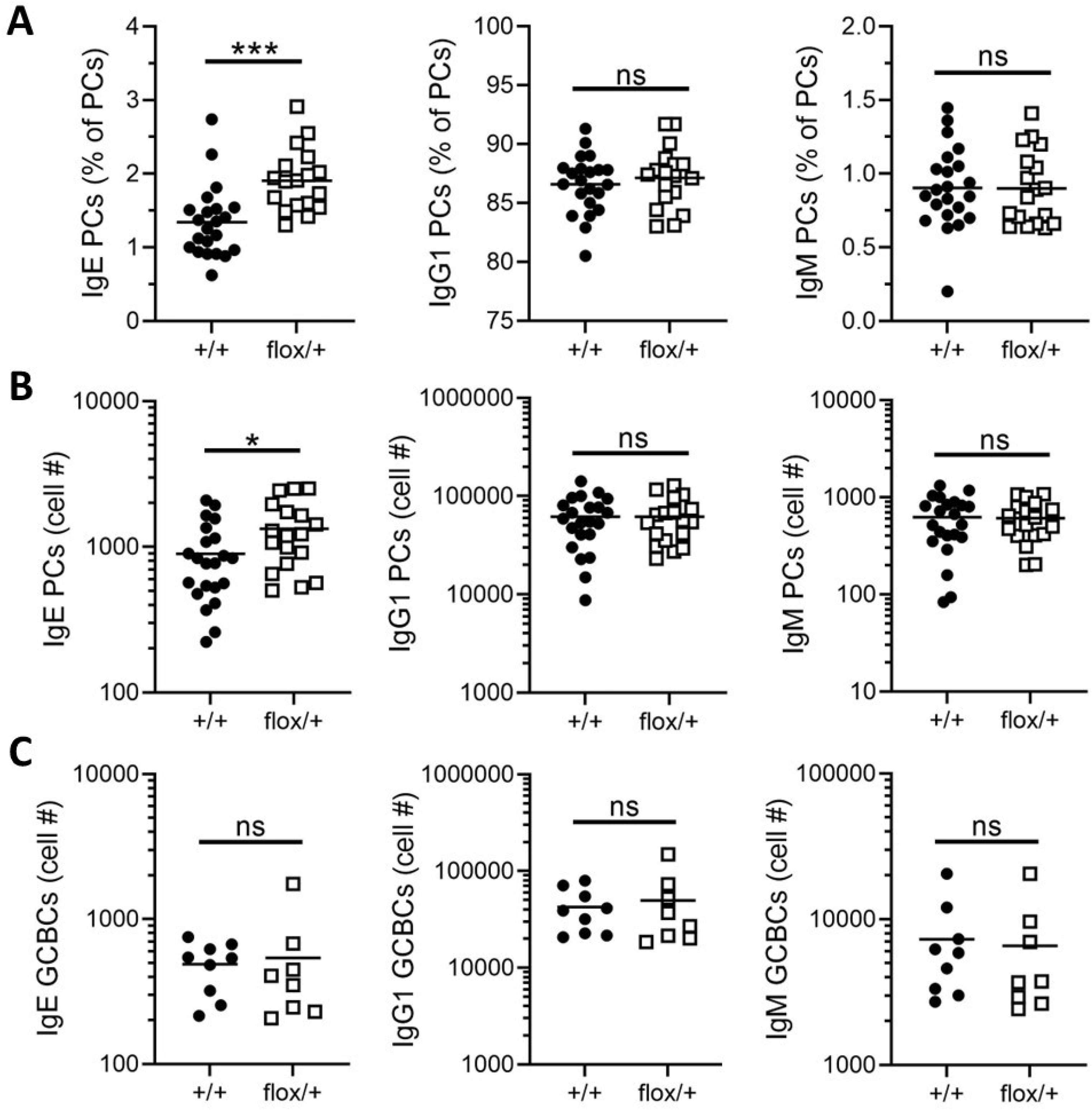
IgE PCs are constrained by BCR signaling *in vivo*. (A-C) Bone marrow chimeras were generated from BLTcre donors that were either *Syk*^flox/+^ or *Syk*^+/+^ and then immunized with NP-CGG in alum adjuvant after reconstitution. Quantification of the frequency **(A)** or number **(B-C)** of the indicated isotypes of PCs or GC B cells (GCBCs) in the dLN as determined by flow cytometry 9 days after immunization. Tamoxifen was administered to all mice 6, 7, and 8 days after immunization (see Methods for details). PCs were gated as B220^int^CD138^+^ and negative for other isotypes, GC B cells were gated as B220^+^CD138^−^PNA^hi^CD38^−^IgD^−^ and negative for other isotypes. Dots represent individual mice and bars represent the mean. n.s., not significant; *, P < 0.05; ***, P < 0.001 (unpaired t tests). Data are pooled from (A-B) or are representative of (C) three independent experiments.

### BCR ligation depletes IgE PCs *in vivo*

Based on our earlier observations that the administration of cognate antigen induced BCR signaling in IgE PCs *in vivo*, we next sought to determine whether this signaling was followed by the depletion of IgE PCs, as we had observed *in vitro*. Indeed, the subcutaneous injection of cognate antigen led to a reduction in the frequency of IgE PCs, but not IgG1 or IgM PCs, that bound cognate antigen (Figure 7A). This was due to a ∼2-fold decrease in the number of antigen-specific IgE PCs, versus no change in the number of IgE PCs that were not antigen-specific (Figure 7B), mirroring our earlier *in vitro* results. We did not observe any change in the number of antigen-specific IgG1 or IgM PCs following antigen injection (Figure 7B). These data extend our *in vitro* results and provide evidence that IgE PCs are uniquely susceptible to depletion by cognate antigen *in vivo*.

**Figure 7.**
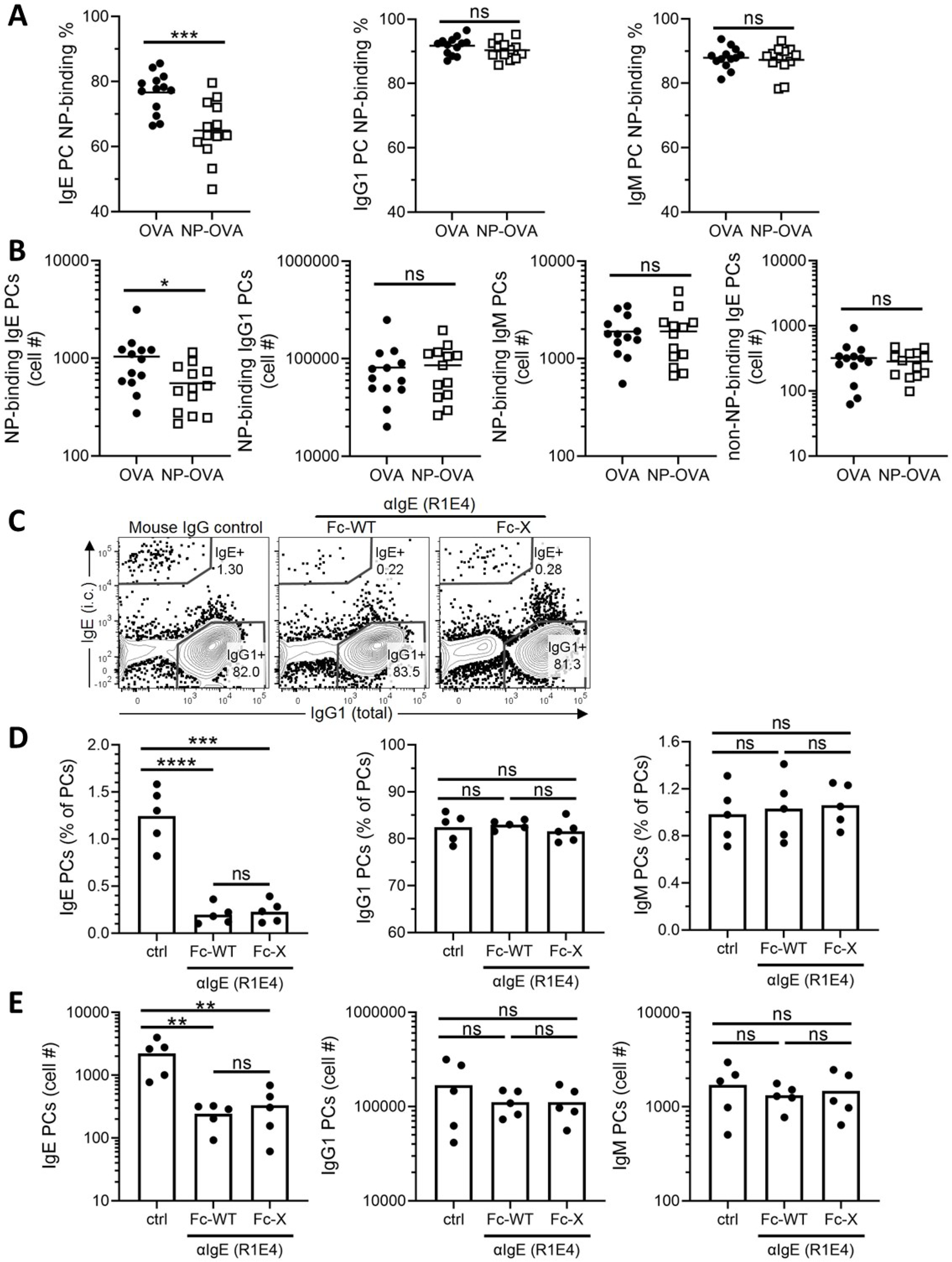
BCR ligation depletes IgE PCs *in vivo*. (A-B) Mice were immunized subcutaneously with NP-CGG in alum adjuvant and injected with control (OVA) or cognate antigen (NP-OVA) 6.33-6.5 days later. The proportion **(A)** and number **(B)** of PCs that were NP-binding were quantified by flow cytometry 16h after antigen injection. The number of non-NP-binding IgE PCs is also shown **(B)**. PCs were gated as CD138^+^B220^int^ and negative for other isotypes. **(C-E)** Mice were immunized with NP-CGG in alum adjuvant and then injected intravenously with αIgE R1E4 (with either a normal Fc region [Fc-WT] or with a mutated Fc region [Fc-X]) or control antibody at 7.33 days after immunization. dLNs were analyzed by flow cytometry 16 hours later. Representative flow cytometry contour plots **(C)** and quantification of the frequency **(D)** and number **(E)** of PCs of the indicated isotypes in the dLN. PCs were gated as CD138^+^B220^int^ and negative for other isotypes. Dots represent individual mice and bars represent the mean. n.s., not significant; *, P < 0.05, **, p <0.01, ***, P < 0.001; ****, P < 0.0001 (unpaired t tests [A-B], one-way unmatched ANOVA with Tukey’s post-test comparing each group to each other group [C-E]). Antigen injection results (A-B) are pooled from two independent experiments while R1E4 Fc-WT versus control comparisons (C-E) are representative of three independent experiments.

To further test if directly ligating mIgE would be a useful strategy to acutely eliminate IgE PC *in vivo*, we took advantage of the unique properties of the R1E4 clone of αIgE that we had previously observed to induce Ca^2+^ flux (Figure S2E) and deplete IgE PCs in cell culture (Figure 4I-J and Figure S4B). This particular αIgE clone only recognizes IgE that is not bound to CD23 or FcεRI, mitigating the possibility that its activity upon IgE PCs could be related to the ligation of receptor-bound IgE (Keegan et al., 1991). Illustrating the potential therapeutic benefit of R1E4, a prior study revealed that recombinant single-chain R1E4 could suppress IgE production in mice (Ota et al., 2009). Therefore, we sought to determine if R1E4 could acutely deplete IgE PCs *in vivo*. WT mice were immunized, and then, after the bulk of IgE PC differentiation had occurred, were administered a single injection of either control antibody, un- modified αIgE R1E4 (Fc-WT), or αIgE R1E4 with a mutated, non-functional Fcγ-receptor binding site (Fc-X). Sixteen hours later, PCs were enumerated in the dLN by flow cytometry.

Injection of αIgE R1E4, but not of control antibody, induced a strong decrease in the frequency (Figure 7C-D) and absolute number (Figure 7E) of IgE PCs. As expected, PCs of other isotypes were unaffected by αIgE R1E4 injection (Figure 7E). The depletion of IgE PCs was equivalent between Fc-WT and Fc-X versions of αIgE R1E4, ruling out a contribution from antibody-dependent cell cytotoxicity (ADCC) or other BCR-independent mechanisms of IgE PC depletion involving Fcγ receptors. These findings demonstrate that ligation of the BCR on IgE PCs *in vivo* results in IgE PC elimination.

## DISCUSSION

Here, we reported that IgE PCs had high surface BCR expression that corresponded with the activation of an intracellular signaling cascade in response to antigen exposure. Specifically, BCR stimulation on IgE PCs resulted in the phosphorylation of Syk, BLNK, and PLCγ2 as well as Ca^2+^ flux. This signaling was toxic to IgE PCs, which were depleted proportionally to the strength of BCR stimulation. Consistent with the induction of apoptosis, BCR stimulation resulted in caspase-3 activation in IgE PCs, which could be rescued from cell death by overexpression of a pro-survival *Bcl2* transgene. Through a combination of pharmacological agents and CRISPR gene targeting, we identified Syk, BLNK, and PLCγ2 as key components of the BCR signalosome required for the elimination of IgE PCs after BCR stimulation. In mice, we found that conditional deletion of a single copy of *Syk* in PCs led to a selective increase in the abundance and relative frequency of IgE PCs, indicating that IgE PCs are uniquely constrained by intrinsic BCR signaling *in vivo*. Conversely, ligation of mIgE *in vivo* with cognate antigen or an αIgE monoclonal antibody led to the elimination of IgE PCs.

Overall, our data have revealed that IgE PCs are uniquely sensitive to BCR signaling-induced elimination. Quantitative differences in BCR signal transduction may explain the increased sensitivity of IgE PCs to antigen-induced elimination compared with IgG1 PCs. We found that IgE PCs had higher surface mIg expression than IgG1 PCs, which correlated with enhanced phosphorylation of BCR signalosome components after the ligation of mIgE compared with mIgG1. Our studies showing that IgE PC elimination was dependent on the affinity, avidity, and amount of antigen binding to the BCR further reinforce this concept. Our findings also imply that qualitative differences in BCR signaling promote the elimination of IgE PCs relative to IgM PCs. Specifically, we found that IgE and IgM PCs had similar surface mIg expression and phosphorylation of proximal BCR signaling proteins, yet IgE PCs were more susceptible to apoptosis induced by BCR stimulation. Furthermore, conditional deletion of *Syk* in PCs *in vivo* led to an increase in IgE PCs but not IgM PCs. The differences in the BCR ligation-induced elimination of IgE and IgM PCs could be related to structural differences between these BCR isotypes. For example, mIgE has an extended cytoplasmic tail (CT), whereas mIgM has a truncated sequence of only three amino acids. The CTs of mIgE and mIgG are thought to prevent recruitment of the inhibitory molecule CD22, which may amplify signaling (Sato et al., 2007). In addition, mIgE and mIgG have a conserved tyrosine motif in their CTs that is thought to be important for amplification of BCR signal transduction (Engels et al., 2009). The mIgE CT has also been reported to associate with HAX-1, which could play a role in the induction of or protection from apoptosis (Laffleur et al., 2015; Oberndorfer et al., 2006). Arguing against a role for the mIgE CT in promoting IgE PC apoptosis, mice with mutations of the tyrosine motif in the mIgE CT, or truncation of the mIgE CT, have diminished, rather than augmented, IgE responses (Achatz et al., 1997; Schmitt et al., 2020). It is unclear whether this result is due to a role for the mIgE CT in IgE PCs or their activated IgE B cell precursors. Overall, further studies will be needed to elucidate the mechanistic basis for the different functional outcomes of ligation of the IgE BCR versus IgG1 and IgM BCRs in PCs.

Relatedly, we found that BCR ligation has different functional consequences in PCs compared with B cells. Our cell sorting experiments showed that BCR ligation induced robust caspase-3 activation in IgE PCs, followed by their depletion, with more modest effects on IgG1 and IgM PCs. However, BCR ligation did not induce aCas3 in activated B cells of any isotype (IgE, IgG1 or IgM) and instead actually increased their abundance. These outcomes may be related, in part, to our culture conditions, which included αCD40 and IL-4 to mimic T cell help. Previous studies have provided conflicting data on whether there are differences in the survival of IgE versus IgG1 B cells in cell culture without BCR stimulation (Haniuda et al., 2016; Laffleur et al., 2015; Newman and Tolar, 2021; Ramadani et al., 2019; Yang et al., 2016), which we have reviewed in detail (Wade-Vallance and Allen, 2021). Our studies reported here are distinct because we specifically investigated the impact of BCR stimulation on cell survival. Overall, our present data indicate that BCR ligation results in signal transduction that is toxic to IgE PCs, but not to IgE B cells.

Our observation that apoptotic PCs lost expression of the marker CD138 also has important implications for the study of apoptosis in the context of IgE responses. Since a large fraction of IgE B cells undergo PC differentiation, apoptotic IgE PCs that have lost CD138 expression could potentially contaminate analyses of IgE B cells. Contamination of dying IgE PCs into IgE B cell flow cytometry gating schemes is especially of concern *in vitro*, where dead cells are not cleared by phagocytes. Therefore, we advise that *in vitro* studies of IgE B cells should take special precautions to exclude dying IgE PCs, which we found have high intracellular IgE staining. Our results further suggest that future studies of PC apoptosis should utilize markers other than CD138 to identify dying PCs.

The studies presented here involving pharmacological inhibitors and CRISPR-Cas9 targeting revealed that Syk, BLNK, and PLCγ2 were essential for the elimination of IgE PCs after BCR stimulation. In contrast, the inhibition of Btk with ibrutinib only partially rescued IgE PCs. This result was unexpected, given the established role of Btk in activating PLCγ2 downstream of BCR stimulation (Rodriguez et al., 2001; Watanabe et al., 2001). Some residual PLCγ2 phosphorylation was reported in a Btk-deficient B cell line, which may be due to the activity of Syk or Src family kinases (Ozdener et al., 2002; Rodriguez et al., 2001; Takata and Kurosaki, 1996). Alternatively, these findings could be explained by redundancy between Btk and other Tec-family kinases for PLCγ2 activation in IgE PCs. A third possibility is that the requirement for PLCγ2 in BCR stimulation-induced IgE PC elimination involves not only its enzymatic effects but also its reported role in stabilizing the assembly of the BCR signalosome (Wang et al., 2014). While the partial contribution of Btk requires further investigation, our data establish a crucial role for the BCR signalosome components Syk, BLNK, and PLCγ2 in the elimination of IgE PCs following BCR stimulation.

The above findings regarding the BCR signalosome provide new insights into prior data obtained from mice with *Syk* and *Blnk* mutations. Specifically, we previously observed that when a single copy of *Syk* was conditionally deleted in activated B cells in mice, this resulted in an increase in the abundance of IgE PCs (Yang et al., 2016). While this could have reflected signaling mediated by Syk in precursor B cells, here we found that conditional deletion of *Syk* in PCs in mice also led to an increase in the abundance of IgE PCs. Together with our cell culture data showing that Syk inhibition prevented IgE PC apoptosis induced by BCR stimulation, our findings provide evidence that *Syk* heterozygosity directly increases the survival of IgE PCs. In BLNK-deficient mice, IgE PCs accumulated for several weeks, suggesting they were long-lived (Haniuda et al., 2016). While this could have been secondary to enhanced IgE GC B cell responses in these mice, our data here indicate that BLNK-deficiency directly protects IgE PCs from apoptosis after BCR stimulation. Therefore, we propose that the increases in IgE PC responses in mice with perturbations in BCR signal transduction are at least in part due to a direct physiological role for BCR signaling in restraining IgE PC survival.

The idea that BCR signaling in PCs impairs longevity is supported by evidence from mice in which the gene encoding SHP-1, a negative regulator of BCR signaling, was conditionally deleted following B cell activation. These studies identified several features of SHP-1-deficiency that seem reminiscent of IgE responses, including increased PC apoptosis, an abortive GC response, and a reduction in long-lived PCs (Li et al., 2014; Yam-Puc et al., 2021). One study also identified a role for SHP-1 in PC migration (Li et al., 2014). Interestingly, prior work swapping the mIgE membrane and cytoplasmic domains with those of IgG1 showed that this also affected bone marrow migration (Achatz-Straussberger et al., 2008). However, in the case of SHP-1 this reportedly affected integrin activity (Li et al., 2014) whereas IgE domain swapping reportedly affected migration to CXCL12 (Achatz-Straussberger et al., 2008). Overall, this evidence is consistent with the model that BCR signaling limits IgE PC longevity *in vivo*, and suggests that this could occur through the dual mechanisms of promoting apoptosis and impairing migration to PC survival niches.

The major finding reported here, that BCR stimulation induced IgE PC elimination, does not exclude a contribution of antigen-independent BCR signaling to the regulation of IgE PC longevity. Elevated BCR expression on IgE PCs may result in increased antigen-independent mIgE signaling relative to IgE B cells, which could negatively impact the survival of IgE PCs. Consistent with this possibility, a recent study reported a higher proportion of IgE PCs than IgG1 PCs had aCas3 staining in an *in vitro* culture system without antigen (Newman and Tolar, 2021). In this study, the authors also noted a reduction in the frequency of aCas3^+^ IgE PCs after CRISPR targeting of *Plcg2* or pharmacological inhibition of Syk. However, this latter finding was not compared to IgG1 PCs. Further exploring how antigen-independent and antigen-dependent BCR signaling govern IgE responses represents an important area of future inquiry, especially *in vivo* where immunization with antigen makes it difficult to disentangle antigen-dependent and antigen-independent effects.

While in this study we focused on the antigen-induced elimination of IgE PCs in mice, we anticipate that these findings will translate to humans. As in mice, human IgE PCs have elevated surface mIgE expression relative to human IgE B cells (Ramadani et al., 2017). Related observations in IgA and IgM PCs are also consistent between mice and humans, including that these cells express elevated mIg and show evidence of BCR signal transduction following mIg ligation (Blanc et al., 2016; Pinto et al., 2013). Similar to our finding that αIgE R1E4 depleted IgE PCs, studies of mice in which a human M1’ region was inserted into mIgE showed that treatment with a therapeutic anti-M1’ antibody led to a reduction in IgE PCs (Brightbill et al., 2010). While the authors of this study attributed this result to effects of the anti-M1’ antibody on IgE B cells that were the precursors of IgE PCs, we propose that this therapy could also directly trigger the apoptosis of IgE PCs. Relatedly, our finding that the R1E4 clone of αIgE eliminated IgE PCs *in vivo* raises the possibility that human therapeutic αIgE monoclonal antibodies, such as omalizumab and legilizumab, could similarly ligate mIg on IgE PCs and induce apoptosis. In this regard, we note that the R1E4 clone of αIgE used in our studies only recognizes IgE that is not bound to FcεRI (Keegan et al., 1991), similar to the omalizumab and legilizumab αIgE therapeutics developed for the treatment of allergic diseases (Gasser et al., 2020). Interestingly, legilizumab was reported to inhibit IgE production in cell culture (Gasser et al., 2020), which we propose could be due to the direct elimination of IgE PCs.

Our conclusion that cognate antigen recognition results in IgE PC elimination predicts that the amount of antigen IgE PCs are exposed to may regulate antigen-specific IgE production. Notably, physiological exposures to inhaled aeroallergens in the respiratory tract may be at relatively low doses compared to those used in mouse models of allergic airway disease. Classic studies found that immunization of rodents with low doses of antigen produced more persistent IgE responses (Holt and McMenamin, 1989; Prouvost-Danon et al., 1977; Vaz et al., 1971).

Powerful IgE responses can also be provoked by anti-IgD immunization in mice (Finkelman et al., 1987), which by definition would be unable to ligate mIgE following IgE class switch recombination. Infections with helminth worms are also known to result in dramatic increases in total IgE, yet most of this IgE response is non-specific (Jarrett et al., 1980). In these infections, high doses of antigen exposure could result in the elimination of antigen-specific IgE PCs, favoring the survival of non-antigen-specific IgE PCs. In human food allergy, patients with severe symptoms typically alter their diet to avoid consumption of the foods to which they are allergic. This allergen avoidance may result in enhanced survival of antigen-specific IgE PCs, consistent with the detection of antigen-specific IgE PCs in the blood of patients with peanut allergy (Croote et al., 2018). Conversely, regular oral exposure to food allergens early in life, such as peanut, has been reported to promote allergen-specific tolerance (du Toit et al., 2018).

We also note that repeated dosing with increasing amounts of antigen is the key concept underlying allergen immunotherapy. Although such regimens likely have many effects on the immune system, including the induction of antigen-specific IgG4 and impacts on T cells (Akdis and Akdis, 2014), our results reveal that the administration of cognate antigen can directly eliminate antigen-specific IgE PCs. This represents a new possible mechanism by which allergen immunotherapy may achieve therapeutic benefit. Overall, we propose that the amount of antigen exposure may be an important factor constraining the lifespan of IgE PCs.

Our findings also raise the possibility that genetic variations in molecules involved in BCR signal transduction or apoptosis could affect IgE PC survival. In support of this idea, a wealth of literature links elevated IgE and/or atopy with primary immunodeficiencies or mutations associated with impairments in antigen receptor signaling pathways (Sokol and Milner, 2018), although aberrations of these pathways would likely affect B cells and T cells in addition to PCs. A recent mutagenesis screen in mice determined that several mutations in genes with a role in BCR signaling, including *Syk* and *Plcg2*, led to increased IgE production (SoRelle et al., 2021). This study estimated that over a third of the human population may be heterozygous carriers for atopy risk alleles that affect BCR signaling or class switch recombination. Even in individuals without inherited alleles that affect BCR signaling or apoptosis, it is possible that sporadic somatic mutations could occur in B cells or PCs that could result in enhanced IgE PC survival in the context of antigen exposure. Our studies have shown that the mutation of even one copy of a gene involved in BCR signal transduction, such as *Syk*, can result in an increase in IgE PC abundance. A potentially important source of somatic mutations in IgE PC precursors could be the GC, in which somatic hypermutation of antibody variable genes occasionally leads to off-target somatic mutations in other regions of the genome, as is well-known from studies of lymphomagenesis (Cyster and Allen, 2019). We therefore propose that inherited and/or somatic mutations that alter BCR signal transduction or apoptosis could result in enhanced survival of antigen-specific IgE PCs in the context of allergy.

The key conclusion of this study is that antigen-driven BCR signaling eliminates IgE PCs, identifying another important way in which the IgE BCR restricts IgE responses. This work establishes a clear functional role for BCR expression on IgE PCs, which has been poorly understood for any PC isotype. Our findings also raise the possibility of other functional outcomes downstream of BCR activation on IgE PCs or other PC isotypes, such as effects on proliferation, cytoskeletal rearrangements, or metabolism. Overall, we believe that our findings may be of significance for understanding immune tolerance to allergens, with potential therapeutic implications.

## MATERIALS AND METHODS

### Mice and bone marrow chimeras

All mice used for experiments in this study were on a C57BL/6 (B6) background (backcrossed ≥10 generations). Mice for experiments were sex and age-matched between groups as much as possible and both male and female mice were used. For *in vivo* experimentation, mice were at least 6 weeks of age and for *in vitro* experimentation donor animals were at least 5 weeks of age.

BLTcre mice (Robinson et al., 2022), *Cd19*^Cre^ mice (Rickert et al., 1997), *EμBcl2-22* (*Bcl2*)-Tg mice (Strasser et al., 1991), *Igh-J^tm1Dh^* mice (Chen et al., 1993), and Verigem mice (Yang et al., 2012) were maintained in our colony on a B6 background. *IgH*^B1-8i^ mice (012642; B6.129P2(C)- *Igh*^tm2Cgn^/J), *IgH*^B1-8hi^ mice (007594; B6.129P2-*Ptprc*^a^*Igh*^tm1Mnz^/J), and *Syk*^flox^ mice (017309; B6.129P2-*Syk*^tm1.2Tara^/J) were purchased from The Jackson Laboratory and then bred in our colony on a B6 background. The following five strains of mice were used as “WT” throughout the manuscript: Boy/J (002014; B6.SJL-*Ptprc*^a^*Pepc*^b^/BoyJ), NCI B6-Ly5.1/Cr (originally 01B96; B6-Ly5.2/Cr, later renamed to B6Ly5.1/Cr), B6/J (000664; C57BL/6J), NCI B6 (556; C57BL/6NCrl), and B6 Thy1.1 (000406; B6.PL-*Thy1^a^*/CyJ). Boy/J, B6/J, and B6 Thy1.1 mice were originally from The Jackson Laboratory and were maintained in our colony, while NCI B6- Ly5.1/Cr and NCI B6 were purchased from Charles River.

Mice were housed in specific-pathogen-free facilities. Mouse work was approved by the Institutional Animal Care and Use Committee (IACUC) of the University of California, San Francisco (UCSF). Bone marrow chimeras were generated as described (Yang et al., 2012). We allowed at least 6 weeks for full reconstitution of the chimeras before immunization.

### Total B cell purification and Igλ B cell enrichment

All *in vitro* culture experiments were performed with B cells purified by negative selection. B cell purification was generally performed as described (Sullivan et al., 2011) but with the following modifications: Media used was DMEM (Fisher Scientific) containing 4.5 g/L of glucose, 2% fetal bovine serum (FBS, Life Technologies), and 10 mM HEPES (Life Technologies). Centrifugation steps were performed at 400g for 5 minutes. ACK lysis was not performed. αCD11c biotin (clone N418, Biolegend) was used at 1μL per 10^8^ cells, and αTER-119 biotin (clone TER-119, Biolegend) was used at 10μL per 10^8^ cells. 100μL of Dynabeads MyOne Streptavidin T1 (Life Technologies) were used per 10^8^ cells. In the case of Igλ B cell enrichments, 10μL of αIgκ biotin (clone RMK-12, Biolegend) were added per 10^8^ cells and 200μL of Dynabeads MyOne Streptavidin T1 were used per 10^8^ cells. After purification, cells were resuspended in complete RPMI (cRPMI), composed of RPMI 1640 without L-glutamine (Fisher Scientific), 10% FBS, 10mM HEPES, 1X Penicillin Streptomycin L-Gluatmine (Fisher Scientific), and 50μM β-mercaptoethanol (Fisher Scientific). Final purity was assessed by flow cytometry. Total B cell purity routinely exceeded 98%. For Igλ-enriched purifications ∼80% of B cells obtained were routinely Igλ^+^.

### Mouse cell culture and *in vitro* BCR stimulation

All cell culture was performed in a humidified incubator with 5% CO2 at 37°C using cRPMI (see above for ingredients) supplemented with 150ng/mL αCD40 (clone FGK-45; Miltenyi Biotec) and 25ng/mL recombinant murine IL-4 (Peprotech) in 96-well Microtest U-bottom plates (BD Falcon) with a volume of 200µl per well. Purified B cells were cultured at a density of 10^4^ cells per well, while Igλ-enriched B cells were cultured at 10^3^ cells per well. Culture duration varied between experiments as described in figure legends. Cells were plated in triplicate for each condition, except for some timecourse experiments where sextuplicates were used. In some cases, CD45-congenic B cells were co-plated to allow the combined assessment of cells from two different mice, such as in comparisons of B1-8 and B1-8hi B cells. Cell counts from such experiments were scaled up two-fold to account for each sample having had half of the normal cell input.

*In vitro* BCR stimulation with cognate antigen was performed by adding NP(4)BSA (Biosearch Technologies), NP(25)BSA (Biosearch Technologies), or BSA (Sigma-Aldrich) as a control directly to the culture supernatant without a media swap. See figure legends for timing and doses.

### Media swap for *in vitro* experiments with anti-BCR antibodies

In some experiments, a media swap was performed to remove sIg from the culture which would otherwise block anti-BCR antibodies. First, the culture supernatant was removed. Then cells were washed once with pre-warmed cRPMI and resuspended in “swap” cRPMI. This “swap” cRPMI was not freshly prepared, but rather was an aliquot of the initial plating media which was maintained in the incubator throughout the culture period. This strategy ensured that the stimulatory environment of the cells did not change pre- and post-swap, obviating the burst of B cell activation that occurs if fresh stimulatory reagents are used as well as the drop off in viability which occurs if costimulatory signals are completely withdrawn. After the media swap, Goat polyclonal F(ab’)2 anti-mouse Igκ (αIgκ; LifeSpan Biosciences), rat anti-mouse IgE clone R1E4, control Chrompure Goat IgG (Jackson ImmunoResearch), or control rat gamma globulin (Jackson ImmunoResearch) was added to achieve a final concentration of 2.5μg/mL.

All inhibitors were diluted in DMSO. The concentration of DMSO in culture never exceeded 0.1% and was typically lower. The specific DMSO vehicle control concentration in each experiment was made to be equivalent to the highest concentration of DMSO in any inhibitor-treated well. Inhibitors were added 5 minutes prior to the addition of stimulating antigen/control. See Table S2 for further details on inhibitors and concentrations used.

### CRISPR of B1-8 cells

CRISPR gene targeting was achieved by electroporation of ribonucleoprotein complexes containing Cas9 and single guide RNAs (sgRNAs), adapted from a method published for human B cells (Wu et al., 2018). Lyophilized modified sgRNAs (Synthego) were designed using Synthego’s sgRNA design tool based on the following criteria in order of importance: 1) cut site within coding region of exon, 2) targeted exon early in coding sequence of protein, 3) cut site early within sequence of exon, 4) high on-target score, 5) target sequence not closely related to off-target sequences. Synthego’s negative control sgRNA sequence was used as the negative control sgRNA. sgRNAs were resuspended in TE buffer (Synthego) at 80μM. See Table S3 for sgRNA sequences.

Igλ-enriched B1-8 cells were cultured for 2-3 days at 4 *10^5^ cells/well in a 12-well tissue culture plate (BD Falcon) and then washed, resuspended in cRPMI and counted using a Coulter Counter Z2 instrument. After all centrifugation steps, 10μL of DNase I was added to the pellet prior to re- suspension. Cells were washed with D-PBS (Fisher Scientific) and then resuspended in 1M buffer (Chicaybam et al., 2013) consisting of RNAse-free H2O (Life technologies), 5mM KCl (Life technologies), 50mM D-Mannitol (Millipore-Sigma), 15mM MgCl2 (Fisher Scientific), and 120mM NaH2PO4/NaHPO4 (Fisher Scientific) at 2.5-10 * 10^6^ cells/mL. 18-20μL of cells (4.5-20 *10^4^ cells) were then plated into 16-well nucleocuvette strips (Lonza). Each well had also previously received 4μL of ribonucleoprotein, which was pre-assembled by gently mixing 2μL 80μM sgRNA with 2μL 40μM Cas9 (UC Berkeley QB3 MacroLab) and incubating at RT for 15- 60 minutes. 1μL of 100μM homology-directed repair template (HDRT) with no homology to the mouse or human genome was added to improve editing efficiency. Cells were then electroporated using an Amaxa 4D nucleofector electroporation with the pulse code EN-138.

Following electroporation, “rescue” cRPMI was added to the electroporated wells and cells were incubated for 15 minutes. Afterwards, 7 *10^3^ – 3 *10^4^ electroporated cells were plated into each well of a previously-prepared culture plate. 2.5-3.5 days following electroporation, cells were stimulated with cognate or control antigen and then analyzed 24 ± 3 hours later by flow cytometry.

### Mouse injections (Adoptive transfer, immunization, antibody treatment, and tamoxifen treatment)

Adoptive transfers were performed by intravenous injection of 5 *10^3^ Igλ-enriched B1-8 B cells into the retro-orbital plexus of anesthetized mice. Mice were immunized the following day. All immunizations were performed using NP-CGG (Biosearch Technologies) resuspended at 1mg/mL in D-PBS and mixed 50:50 volumetrically with Alhydrogel (Accurate Chemical and Scientific). Mice were injected subcutaneously with 20μL of this solution (10μg of NP-CGG per injection) in each ear. At endpoint, the facial LNs from each side were pooled for analysis (see figure legends for various timepoints).

Unmodified αIgE (clone R1E4; grown from hybridoma), αIgE with a mutated Fc-receptor binding domain (clone R1E4; Cedarlane), or control rat gamma globulin were diluted in D-PBS to a concentration of 0.3mg/mL and injected intravenously to achieve a final dose of 3.25mg/kg. For the experiment shown, mouse gamma globulin (Jackson ImmunoResearch) was used as a control whereas in previous experiments that the presented data is representative of rat gamma globulin was used as a control.

Tamoxifen was dissolved at 50mg/mL in corn oil (Sigma-Aldrich) by shaking at 56°C for several hours. ∼100μL/mouse was delivered by intraperitoneal injection to achieve a dose of 200mg/kg.

### Production of αIgE clone R1E4

The R1E4 hybridoma was cultured in CELLine bioreactor flasks (Fisher Scientific) in DMEM with Glutamax, 100U/mL Penicillin and 100ug/mL streptomycin, and 55mM β-Mercaptoethanol). The hybridoma was seeded at 10^6^ cells/mL in 15 mL 20% FBS in cDMEM, and the nutrient chamber filled with 500-1000 mL 10% FBS in cDMEM. The culture was harvested after seven days and the supernatant purified into PBS by Protein G affinity using a HiTrap Protein G HP affinity column (GE Healthcare), eluted with 0.1M acetic acid (VWR international) pH=3.0 on an AktaPure chromatography system. The eluate was neutralized with 1/10^th^ volume of 1.5M NaCl, 1M Tris-HCL pH=8.0 (Fisher Scientific). Fractions containing protein as determined by the A280 absorbance were combined and dialyzed into PBS with 10k MWCO Slide-A-Lyzer dialysis cassettes (Fisher Scientific). The dialysed antibody was concentrated in a 50 MWCO Amicon Ultra Spin column (Fisher Scientific), aliquoted in PBS at 2.5mg/mL, and then frozen at -80°C until use.

### Flow cytometry

For analysis of *in vivo* experiments, cell suspensions were prepared from dLNs as described (Yang et al., 2012), counted using a Coulter Counter Z2 instrument, and then plated in triplicate at a density of 3.5 * 10^6^ cells per staining well. Cells were then stained with antibodies (Table S1) essentially as described (Yang et al., 2012), with one notable difference being that sodium azide was excluded from FACS buffer (2% FBS and 1mM EDTA [Fisher Scientific] in PBS) to ensure optimal CD138 staining (Wilmore et al., 2017). All incubations were 20 minutes on ice except for Fc block incubations (10 minutes) and antibody staining of fixed and permeabilized cells (45 minutes to 1 hour). After the final staining centrifugation step was complete, cells from triplicate staining wells were resuspended and pooled using 100µL FACS buffer and then combined with 7500 AccuCount Ultra Rainbow Fluorescent counting beads (Spherotech; hereafter referred to as counting beads) per sample prior to analysis.

At the endpoint of *in vitro* experiments, supernatants were removed by pipetting, triplicate culture wells (or sextuplicate wells, for some timecourse experiments) were pooled using 100µL FACS buffer per sample, and then the solution of resuspended cells was transferred to a staining plate previously prepared with 7500 counting beads per well. After pelleting and removal of the supernatant, staining was performed as described above for *in vivo* experiments. Cells were then resuspended in 25µL FACS buffer prior to analysis.

For both *in vivo* and *in vitro* experiments, we used our previously-established intracellular staining technique (Yang et al., 2012) to sensitively and specifically detect IgE-expressing cells. Briefly, to prevent the detection of IgE captured by non-IgE-expressing cells, surface IgE was blocked with a large excess of unconjugated αIgE (clone RME-1). IgE-expressing cells were then detected after fixation/permeabilization by staining with a low concentration of fluorescently-labelled RME-1.

After staining, cells were collected on an LSRFortessa (BD). Data were analyzed using FlowJo v10. Counting beads were identified by their high SSC and extreme fluorescence and were used to determine the proportion of the cells plated for staining that had been collected on the flow cytometer for each sample. Cells were gated on FSC-W versus FSC-H and then SSC-W versus SSC-H gates to exclude doublets. Except where otherwise specified, cells were also gated as negative for the fixable viability dye eFluor780 and over a broad range of FSC-A to capture resting and blasting lymphocytes. 2D plots were presented as either dot plots or contour plots with outliers shown as dots. In some cases, ‘large dots’ were used to visualize rare events.

### *In vivo* NP-OVA injection experiments for phosflow and PC depletion assays

Mice were immunized as described above. At D6.33-D6.5, mice were administered 200µg of cognate antigen (NP conjugated to ovalbumin [NP(16)OVA, Biosearch Technologies]) or control antigen (ovalbumin; OVA [Worthington Biochemicals]) in 20µl PBS by subcutaneous injection, such that the injected antigen would drain to the same LN(s) as the initial immunization. To assess PC depletion, dLNs were analyzed by flow cytometry 16h later. For phosflow analysis of proteins involved in proximal BCR signal transduction, mouse euthanasia was timed so that dLNs could be put in ice-cold PBS with eFluor780 (1/3000) 15-30 min after antigen injection. dLNs were processed into a single cell suspension on ice, combined with an equal volume of 4% PFA, and then incubated for 10 minutes in a 37°C water bath to fix the cells and stop BCR signaling. Fixed cells were washed and then a surface stain was performed as above. Afterwards, cells were incubated with 1.6% PFA for 5 minutes at room temperature and then centrifuged at 1000 × *g*. The supernatant was discarded and then cells were incubated in the residual volume on ice for 5 minutes. Next, cells were permeabilized by adding three consecutive 30µL increments of ice-cold 90% methanol in PBS and mixing with a pipette. Cells were incubated on ice for 30 minutes, then 110µL FACS buffer was added, and cells were incubated for a further 5 minutes. Cells were then washed twice by centrifugation at 1000 × *g*, resuspended in 200µL FACS buffer, and centrifuged again. Washed cells were then incubated with anti-isotype antibodies for 45 minutes on ice. Afterwards, cells were washed twice as before, resuspended in 20µL of FACS buffer containing 1-2% normal mouse serum (Jackson ImmunoResearch) and 1-2% mouse gamma globulin, and incubated at RT for 30 minutes. This step occludes the free arms of the anti-mouse IgG1 antibody, which was used to detect IgG1 PCs, to block subsequent binding to phosflow antibodies of the mouse IgG1 isotype. After 30 minutes of blocking, phosflow antibodies were added and cells were incubated for a further 40 minutes, and then cells were washed prior to analysis. See Table S1 for antibodies and dilution factors.

### *In vitro* phosflow experiments

A media swap was performed as described above and then cultured cells were incubated at 37°C for 5 minutes with 12.5µg/mL goat anti-mouse Igκ F(ab)’2 and eFluor780 (1/1000). Cells were then fixed by addition of PFA to a final concentration of 2% and incubated at 37°C for 10 minutes to stop BCR signaling and fix the cells. The rest of the protocol was performed as described above for *in vivo* phosflow.

### Ca^2+^ flux experiments

All Ca^2^ flux experiments were performed with cells from mice compound heterozygous for the Verigem and *Igh-J^tm1Dh^* IgH alleles. This strategy ensured that mIgE^+^ BCR expression originated from the Verigem allele, such that all IgE^+^ cells were Venus^+^. B cells were purified and cultured as described above for 4.5 days ± 6 hours. Cells were then washed and resuspended in 100μL PBS + 1% FBS. Each time cells were centrifuged, 10μL DNase I (Roche) was added to the pellet prior to re-suspension. Cells were then counted with a Coulter Counter Z2 instrument, diluted to 20 * 10^6^ cells/mL, and stained with eFluor780 Fixable Viability Dye (1/1000) and CD138-PE (1/200). Cells were stained for 5-15 minutes at RT or 37°C. Cells were washed, resuspended in cRPMI at 20 * 10^6^ cells/mL, and loaded with the Ca^2+^-sensitive dye Indo-1 at 0.1μM for 20 minutes at 37°C. Finally, cells were washed again and resuspended in cRPMI at 20-100 * 10^6^ cells/mL and maintained in a water bath at 37°C prior to flow cytometric analysis. Events were collected on an LSR Fortessa X-20 (BD) for ∼10s prior to the addition of anti-BCR antibodies (αIgE clone R1E4 or αIgκ) or control (Goat IgG) at a final concentration of 100μg/mL. After the addition of anti-BCR antibodies, cells were vortexed to mix and returned to the cytometer for observation of Ca^2+^ flux.

### Cell sorting experiments

Cell sorting experiments were performed using Verigem-homozygous cells cultured for 4.5 days ± 6 hours. Cells were retrieved from culture, washed, and resuspended in Hank’s Balanced Salt Solution (HBSS; obtained from UCSF cell culture facility) supplemented with 1% FBS, 0.5% BSA (Sigma-Aldrich), 2mM EDTA and 10mM HEPES, hereafter known as ‘sorting media’.

Cells were then counted with a Coulter Counter Z2 instrument, diluted to 20M cells/mL, and stained with eFluor780 (1/1000), CD138-PE, and IgD-BV510 for 10-30 minutes at RT. Cells were washed, resuspended in HBSS, and diluted to a final concentration of 30 * 10^6^ cells/mL in sorting media. Cells were then sorted on a BD FACS Aria II instrument.

Post-sort media was prepared which consisted of 50% FBS 50% cRPMI. Immediately prior to sorting, post-sort media was passed through a 40μm Falcon cell strainer (Fisher Scientific) into a 5mL polypropylene tube (Fisher Scientific). Cells were sorted with a 70μm nozzle at a frequency of 87,000 droplets/second. Immediately following sorting, the purity of sorted populations was verified by flow cytometry. Sorted cells were plated into previously prepared culture plates at 0.5-2 *10^4^ sorted cells/well. Cells were then treated with anti-BCR antibodies or controls (5μg/mL) as well as with or without various inhibitors and controls and maintained in culture for an additional 24 ± 3 hours prior to flow cytometric analysis.

### Statistical analysis

To achieve power to discern meaningful differences, experiments were performed with multiple biological replicates and/or multiple times, see figure legends. The number of samples chosen for each comparison was determined based on past similar experiments to gauge the expected magnitude of differences. GraphPad Prism v9 was used for statistical analyses. Data approximated a log-normal distribution and thus were log transformed for statistical tests.

Statistical tests were selected by consulting the GraphPad Statistics Guide according to experimental design. All tests were two-tailed.

## SUPPLEMENTARY TABLES

**Table S1:** Antibody-fluorochrome conjugates and other reagents used for flow cytometry

| Antibody target or reagent designation | Clone | Company | Conjugate | Dilution |
| --- | --- | --- | --- | --- |
| Active Caspase-3 | C92-605 | BD Biosciences | V450 | 1/50-100 |
|  |  |  | FITC | 1/25 |
|  |  |  | Alexa Fluor 647 | 1/50-100 |
| B220 | RA3-6B2 | BD Biosciences | V500 | 1/100-200 |
|  |  | Life Technologies | Qdot 655 | 1/100-400 (varies by lot) |
|  |  |  | APC | 1/150-250 |
| CD38 | 90 | Life Technologies | Alexa Fluor 700 | 1/100 |
|  |  | BD Biosciences | Brilliant Violet 786 | 1/200 |
| CD45.1 | A20 | Biolegend | Brilliant Violet 650 | 1/150-200 |
|  |  | BD Biosciences | Alexa Fluor 647 | 1/150-200 |
|  |  |  | Alexa Fluor 700 | 1/100 |
| CD45.2 | 104 | Biolegend | FITC | 1/100-200 |
|  |  |  | PerCP-Cy5.5 | 1/100-200 |
|  |  |  | Brilliant Violet 785 | 1/100 |
|  |  | BD Biosciences | PerCP-Cy5.5 | 1/100-200 |
|  |  |  | PE-Cy7 | 1/100-200 |
| CD138 | 281-2 | BD Biosciences | PE | 1/200 |
|  |  |  | Brilliant Violet 711 | 1/150-200 |
| Fixable Viability Dye eFluor 780 | N/A | Life Technologies | N/A | 1/600-1000 |
| IgD | 11-26c.2a | Biolegend | Brilliant Violet 510 | 1/100-1/200 |
|  |  |  | Brilliant Violet 711 | 1/200 |
|  |  |  | FITC | 1/200 |
|  |  |  | PerCP-Cy5.5 | 1/200 |
|  |  |  | Alexa Fluor 700 | 1/100 |
| IgE | RME1 | Biolegend | Unconjugated | 1/15 |
|  |  |  | FITC | 1/400-500 |
|  |  |  | PE | 1/350-500 |
|  | R35-72 | BD Biosciences | Unconjugated | 1/15 |
|  |  |  | Brilliant Violet 650 | 1/150-300 |
| IgG1 | A85-1 | BD Biosciences | V450 | 1/300-400 |
|  |  |  | FITC | 1/300-400 |
| Ig$\lambda_1$ | R11-153 | BD Biosciences | Biotin | 1/300-1/400 |
| Ig$\lambda_{1,2,3}$ | R26-46 | BD Biosciences | FITC | 1/150-200 |
| IgM | II/41 | Life Technologies | PE-Cy7 | 1/175-1/500 |
| NP | N/A | Conjugated in-house as described (Yang et al., 2016). | APC | 1/750 of 0.1mg/mL stock |
| PNA (peanut agglutinin) | N/A | Vector Laboratories | Biotin | 1/1000-2000 |
|  |  |  | FITC | 1/500 |
| p-Syk (pY348) | I120-722 | BD Biosciences | Alexa Fluor 488 | 1/5 |
| p-BLNK (pY84) | J117-1278 | BD Biosciences | Alexa Fluor 647 | 1/5 |
| p-PLCγ2 (pY759) | K86-689.37 | BD Biosciences | Alexa Fluor 488 | 1/5 |
| Streptavidin | N/A | Life Technologies | QDot 605 | 1/400 |
|  |  | BD Biosciences | Brilliant Violet 711 | 1/400 |
| TruStain FcX (anti-mouse CD16/32) | 93 | Biolegend | Unconjugated | 1/50 |

**Table S2:** Inhibitors

| Inhibitor name | Company | Final concentration |  |  |
| --- | --- | --- | --- | --- |
|  |  | “Low” | “Medium” | “High” |
| Ibrutinib | Gift from Dr. Jack Taunton | 500pM | 2nM | 20nM |
| PRT062607 | Selleck Chemicals | 250nM | 1μM | 4μM |

**Table S3:**
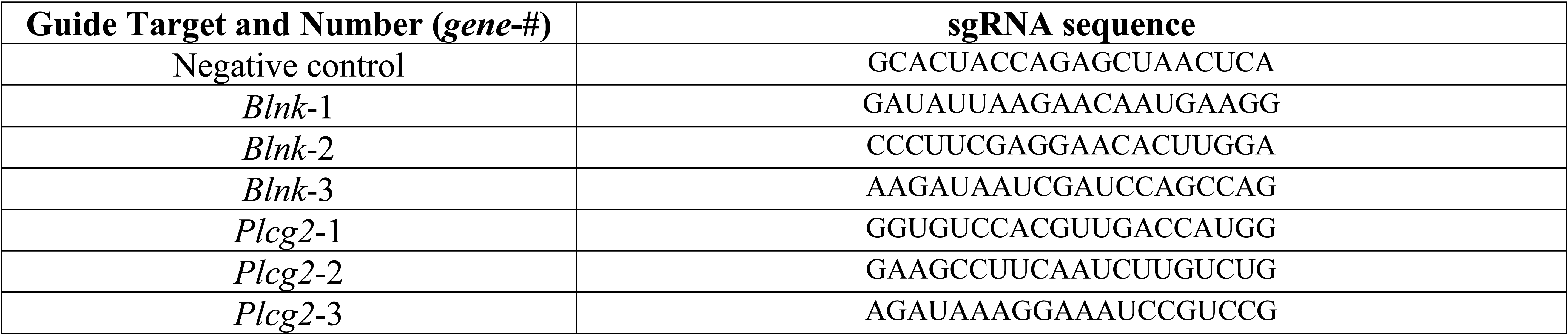
sgRNA sequences

## Acknowledgments

We thank J.B. Jung for assistance with bone marrow chimera experiments; J. Taunton and G.A. Smith for generously providing ibrutinib for our studies; S. Elmes from the Laboratory for Cellular Analysis Core (supported by NIH Grant P30CA082103) for technical assistance with cell sorting experiments; the Gladstone Flow Cytometry Core (supported by NIH Grant P30 AI027763) for their technical support with Ca^2+^ flux experiments; and J.G. Cyster, K.M. Ansel, J. Zikherman, and J.F.E. Koenig for advice and comments on the manuscript.

## Author contributions

A.K.W-V., Z.Y., and C.D.C.A. designed the experiments. Z.Y. performed initial phosflow and apoptosis experiments. J.B.L. performed most inhibitor experiments and assisted with several other experiments. A.K.W-V performed the remainder of the experiments. M.J.R. characterized the BLTcre mice and grew the R1E4 hybridoma. D.M.T. supervised the generation of the BLTcre mice. C.D.C.A. supervised the research. A.K.W-V. and C.D.C.A. analyzed the data and wrote the paper.

## Funding

This research was supported by the National Institute of Allergy and Infections Diseases of the National Institutes of Health under Award Number R01AI130470; the Program for Breakthrough Biomedical Research, which is partially funded by the Sandler Foundation; the Sandler Asthma Basic Research Center; and the Cardiovascular Research Institute at the University of California, San Francisco. C.D.C.A. was a Pew Scholar in the Biomedical Sciences, supported by The Pew Charitable Trusts. A.K.W-V was supported by a Doctoral Foreign Study Award from the Canadian Institute of Health Research, funding reference number DFD-170769. D.M.T. and M.J.R. were funded by National Health and Medical Research Council (NHMRC) Australia by an Investigator Award (APP1175411) and Ideas Grant (APP1185294), respectively. The content is solely the responsibility of the authors and does not necessarily represent the official views of the funding agencies.

## Conflict of interest statement

C.D.C.A. is a member of the scientific advisory board for Walking Fish Therapeutics. C.D.C.A.’s spouse is an employee and shareholder of Bristol Myers Squibb. Z.Y. is an employee of Genentech and a shareholder in the Roche Group. These companies had no involvement in this work. The authors have no other financial conflicts of interest to declare.

---

**Figure S1.**
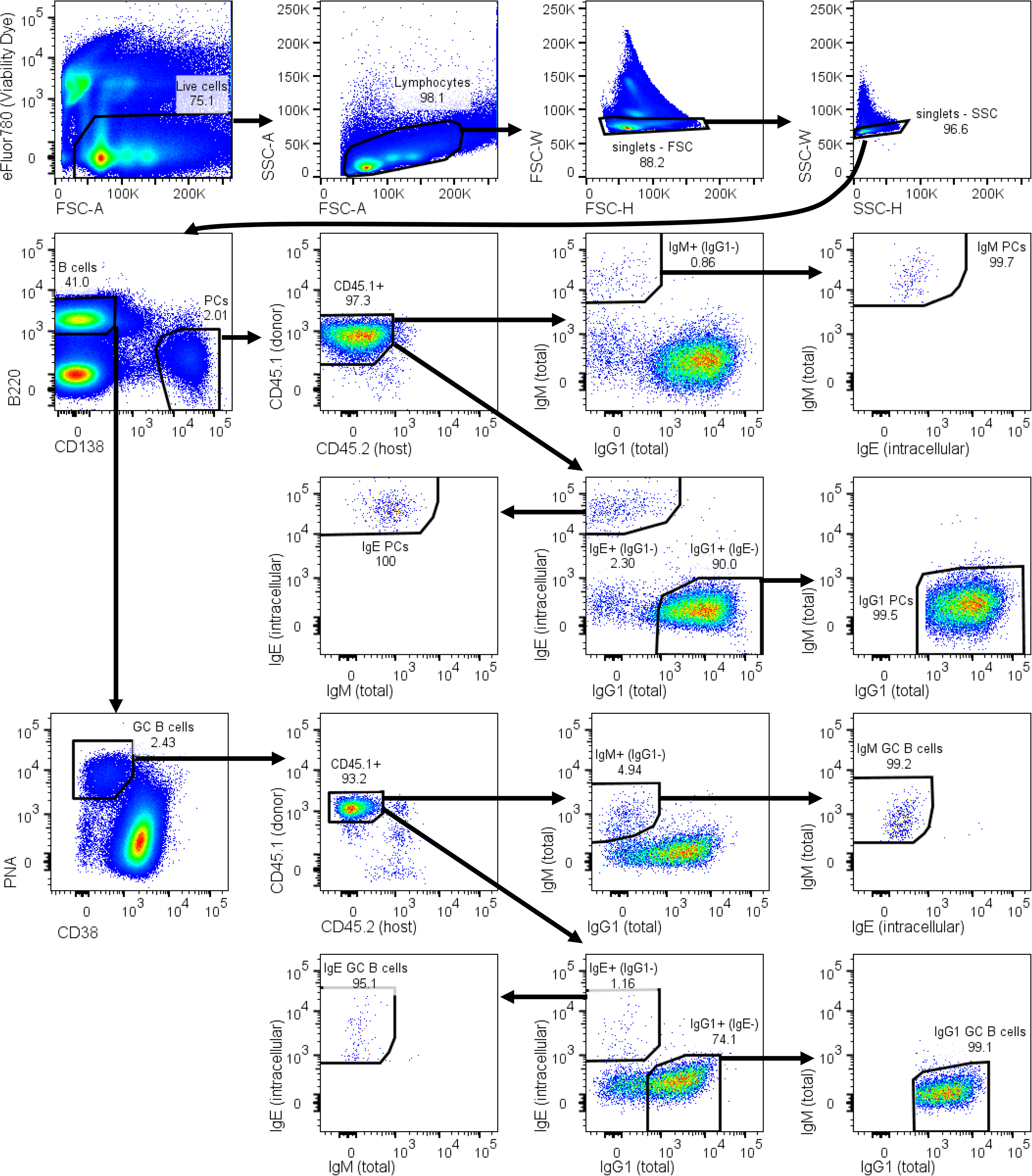
Representative flow cytometric analysis of cells from draining LNs. Cells were gated for viability, as lymphocytes based on their scatter characteristics, and then as singlets by pulse width versus height. Cells were then sub-divided into B cell and PC populations. In this particular representative gating strategy, cells were also gated on congenic markers (CD45.1^+^ CD45.2^−^) to isolate transferred cells. PCs were then gated as highly expressing intracellular Ig of the indicated isotype without expression of Ig of any other isotype by sequential gating. GC B cells were gated as PNA^hi^CD38^+^ and then by isotype, using a similar strategy as for PCs. All plots shown are from the same representative sample.

**Figure S2.**
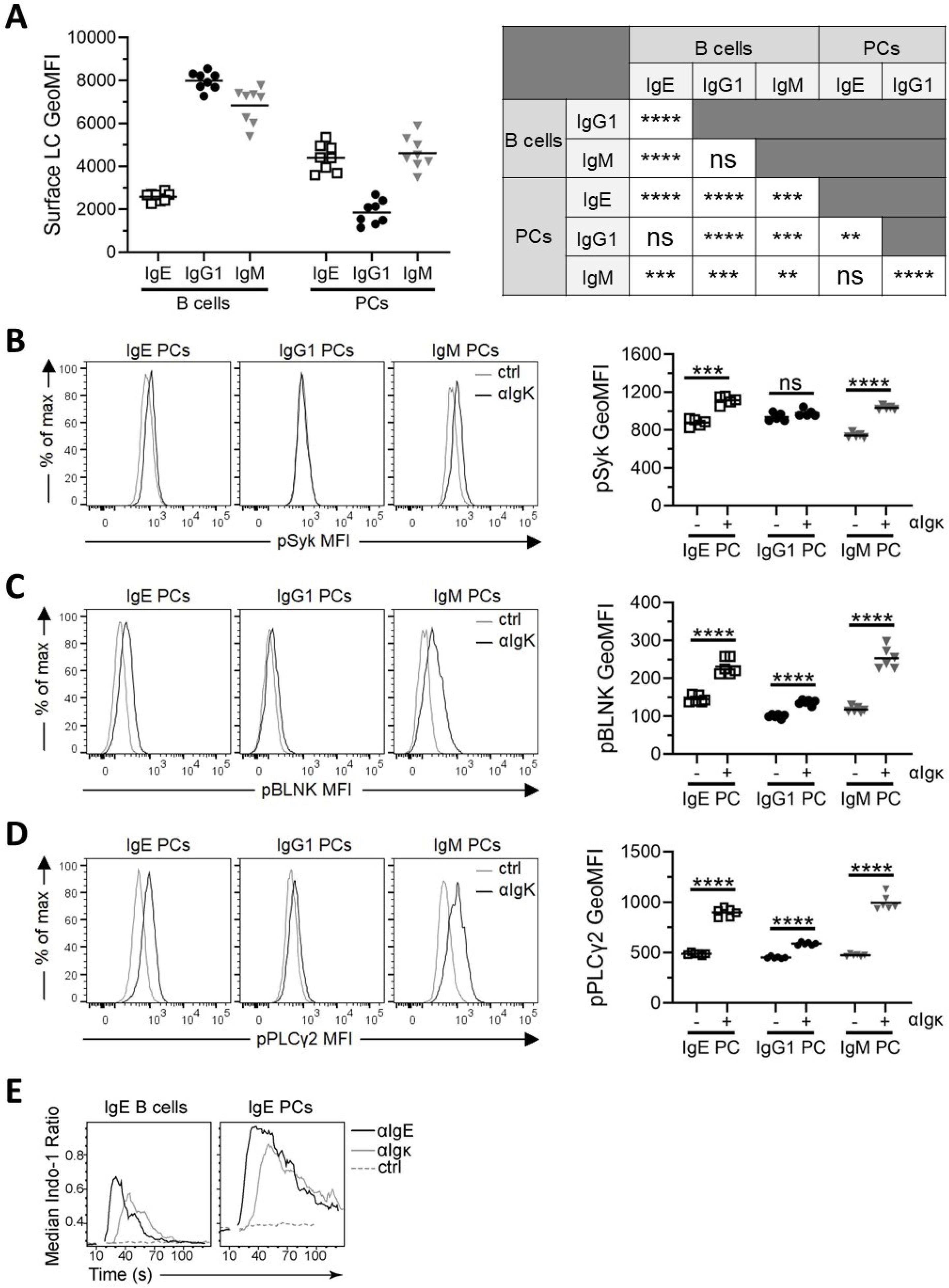
**(A)** Quantification of the surface Igλ LC expression on the indicated cell types by flow cytometry of Igλ- enriched B1-8 B cells cultured for 5.5 days ± 6h. B cells were gated as B220^+^CD138^−^IgD^−^ and PCs were gated as B220^int^CD138^+^. Residual Igκ^+^ B cells were excluded by gating on NP-binding cells. GeoMFI, geometric mean fluorescence intensity. **(B-D)** Phosflow analysis of pSyk **(B)**, pBLNK **(C)**, and pPLCγ2 **(D)** with representative histograms (left) and quantification (right). Purified B cells were cultured for 4.5 days then treated for 5 minutes with either polyclonal goat anti-Igκ F(ab)’2 antibody (αIgκ; ‘+’) or goat IgG control (‘-’). PCs were gated as B220^int^IgD^−^CD138^+^ and negative for other PC isotypes. **(E)** Purified Verigem B cells were cultured for 4.5 days, after which cells were stained and loaded with Indo-1. Shown are plots of the median ratio of bound:unbound Indo-1 in IgE B cells (left; identified as Venus^lo^CD138^−^) and IgE PCs (right; identified as Venus^hi^CD138^+^) prior to and following administration of the indicated control or stimulating antibodies. Dots represent cells cultured from individual mice and bars represent the mean (A-D). n.s., not significant; **, P < 0.01; ***, P<0.001, ****, P < 0.0001 (one-way ANOVA with Tukey’s post-test comparing the mean of each group with the mean of every other group [A, table], one-way ANOVA with Sidak’s post-test comparing the means of pre-selected pairs of groups [B- D]). Data are pooled from (quantification graphs and table) or representative of (histograms) two (A-D) or three (E) independent experiments.

**Figure S3.**
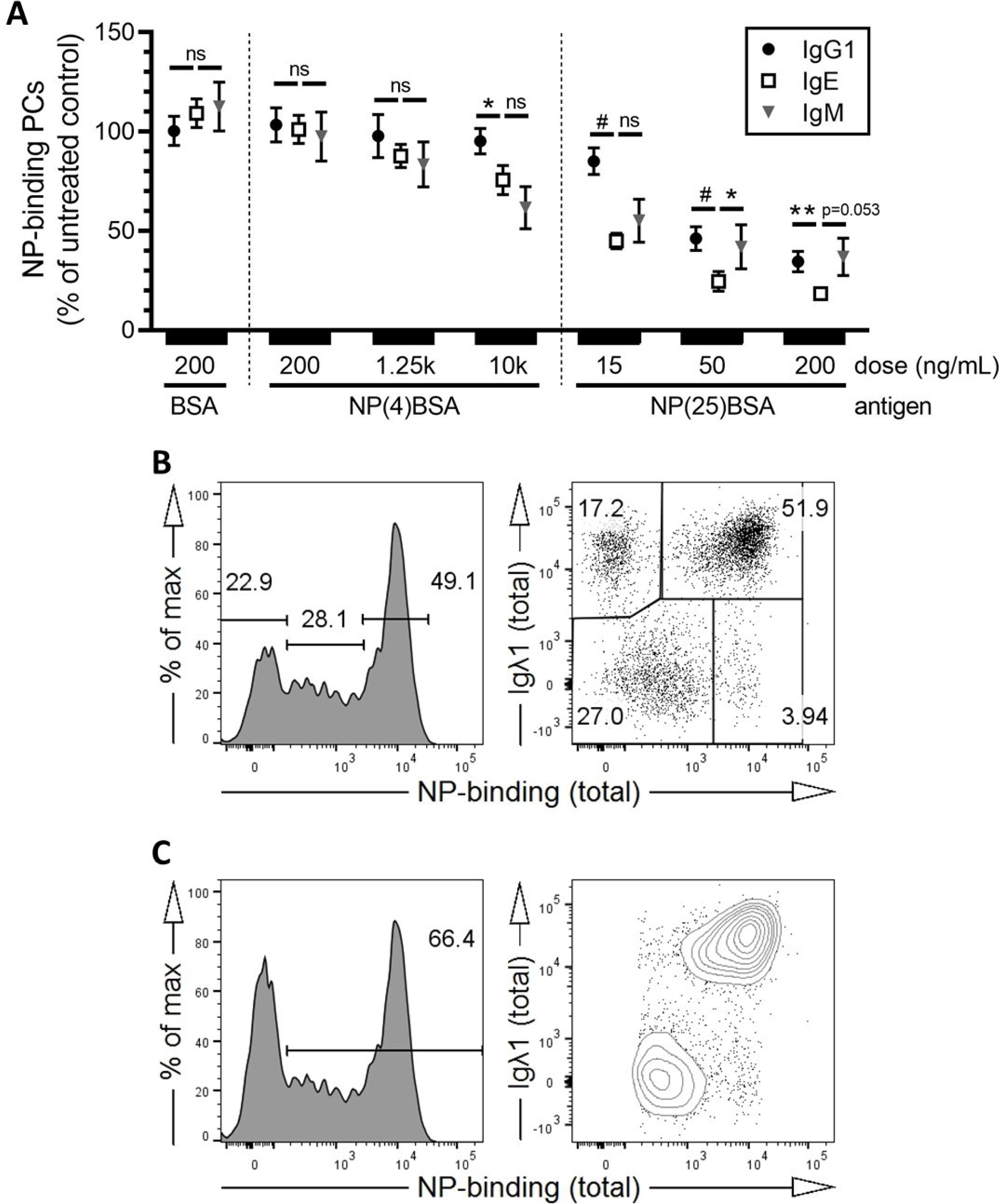
(A-C) Igλ-enriched B1-8 B cells were cultured for 5.25 days ± 7 hours and then analyzed by flow cytometry. **(A)** After treatment with the indicated antigens for the final 12 ± 1 hours of the culture, the number of NP-binding PCs of each isotype was quantified by flow cytometry and is shown as a percentage of the number of NP-binding PCs of the same isotype in untreated wells. Dots represent the mean and error bars reflect the standard error, cells from n=14 mice. **(B)** Left, representative flow cytometry histogram of total (surface + intracellular) NP- APC staining of Igλ-expressing IgE PC divided into non-NP-binding, intermediate NP-binding, and high-NP binding gates. Right, representative flow cytometry dot plot of the same cells comparing NP-binding to Igλ1 staining. **(C)** Left, representative flow cytometry histogram of total NP-APC staining of IgE PC divided into NP- binding (intermediate NP-binding + high NP-binding, as depicted in the left panel of B) and non-NP-binding. Right, representative flow cytometry contour plot of IgE PCs gated as NP-binding (as shown in the left panel). (A-C) IgE PCs were gated as B220^int^CD138^+^IgE^+^. n.s., not significant; *, P < 0.05; **, P < 0.01; #, P < 0.0001 (two-way ANOVA with Dunnett’s post-test comparing IgG1 and IgM to IgE [A]). Data are pooled from (A) or are representative of (B-C) three independent experiments.

**Figure S4.**
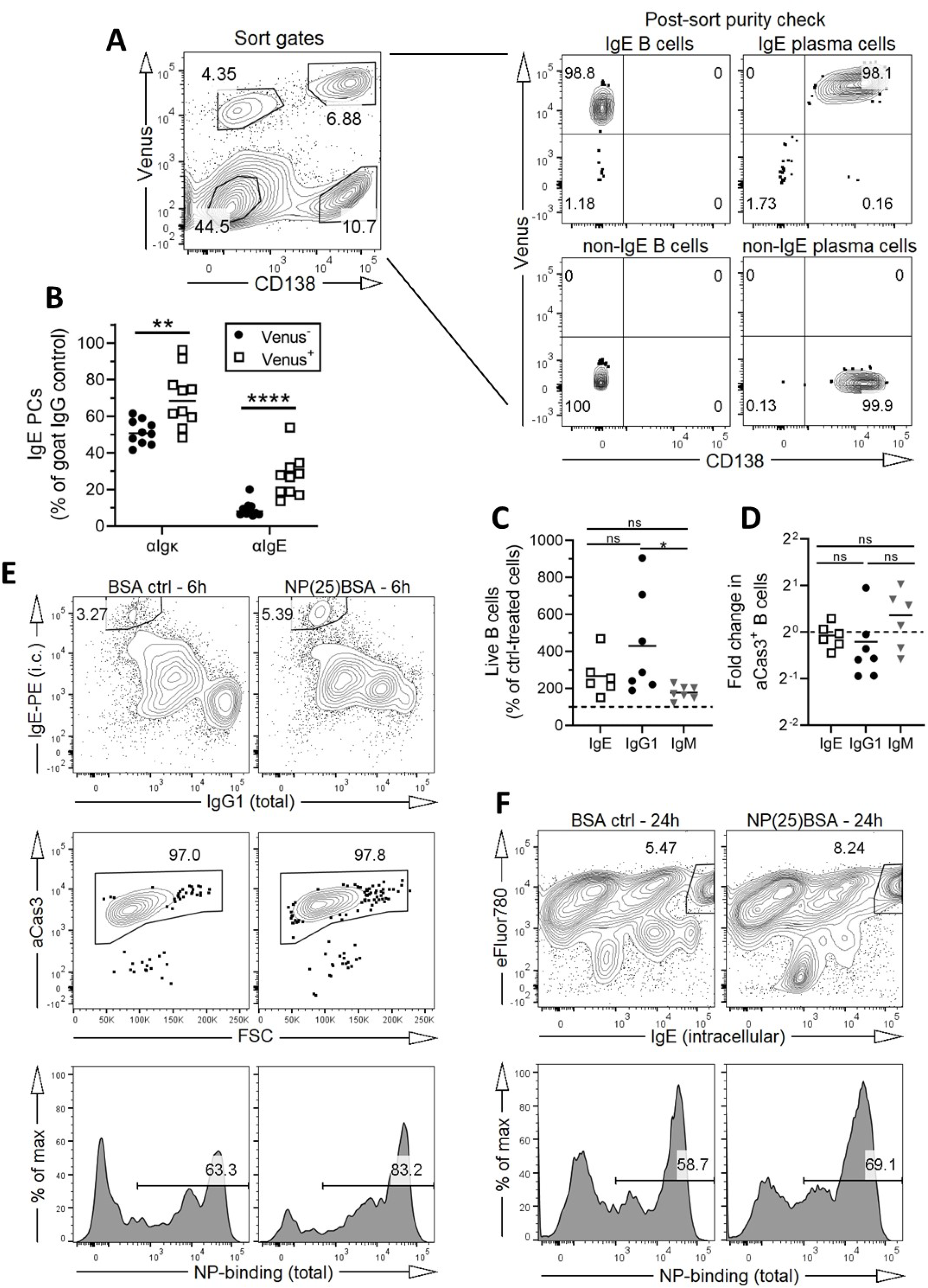
**(A)** For the experiments shown in Figure 4A-C, representative flow cytometry contour plots depict the gates used for cell sorting (left) and a confirmation of the purity of sorted cells (right). **(B)** B cells from Verigem heterozygous mice were purified and cultured for 5.5 days ± 6 hours and then analyzed by flow cytometry. 24 ± 3 hours prior to analysis cells were treated with goat IgG control, αIgκ, or αIgE. The relative number of IgE PCs (B220^int^CD138^+^IgE^+^) after BCR stimulation is shown as a percentage of the number of IgE PCs after treatment with control IgG with the data segregated by whether the cells expressed the Verigem allele (Venus^+^) or the WT allele (Venus^-^). **(C-D)** Cells were sorted by flow cytometry as shown in (A) and cultured as described in Figure 4A-C and Methods. Sorted cells were cultured separately and treated with either goat polyclonal IgG control or αIgκ antibody for 24 ± 3 hours. **(C)** Quantification of the number of B cells after αIgκ treatment shown as a percentage of the number of B cells after control IgG treatment for each isotype. The dotted line represents no change from ctrl. **(D)** The fold change in the proportion B cells that were aCas3^+^ after αIgκ treatment compared to control IgG treatment. The dotted line represents no change from ctrl. **(E)** Representative flow cytometric analysis of cells from the timecourse analysis in Figure 4F at 6 hours. Cells were gated as CD138^−^ and then sequentially from top to bottom showing the identification of IgE^HI^NP^+^aCas3^+^ cells. **(F)** Representative flow cytometric analysis of cells from the timecourse analysis in Figure 4G at 24 hours. Cells were gated sequentially from top to bottom showing the identification of e780^+^IgE^HI^NP^+^ cells. Dots represent data points derived from cells from individual mice. n.s., not significant; *, P < 0.05; **, P < 0.01; ****, P < 0.0001 (paired t tests with the Holm-Sidak correction for multiple comparisons [A], one-way unmatched ANOVA with Tukey’s post-test comparing each group to each other group [C, D]). Bars represent the mean. Results are representative of three (A) or two (E-F) independent experiments or are pooled from three (B-C) or two (D) independent experiments.

**Figure S5.**
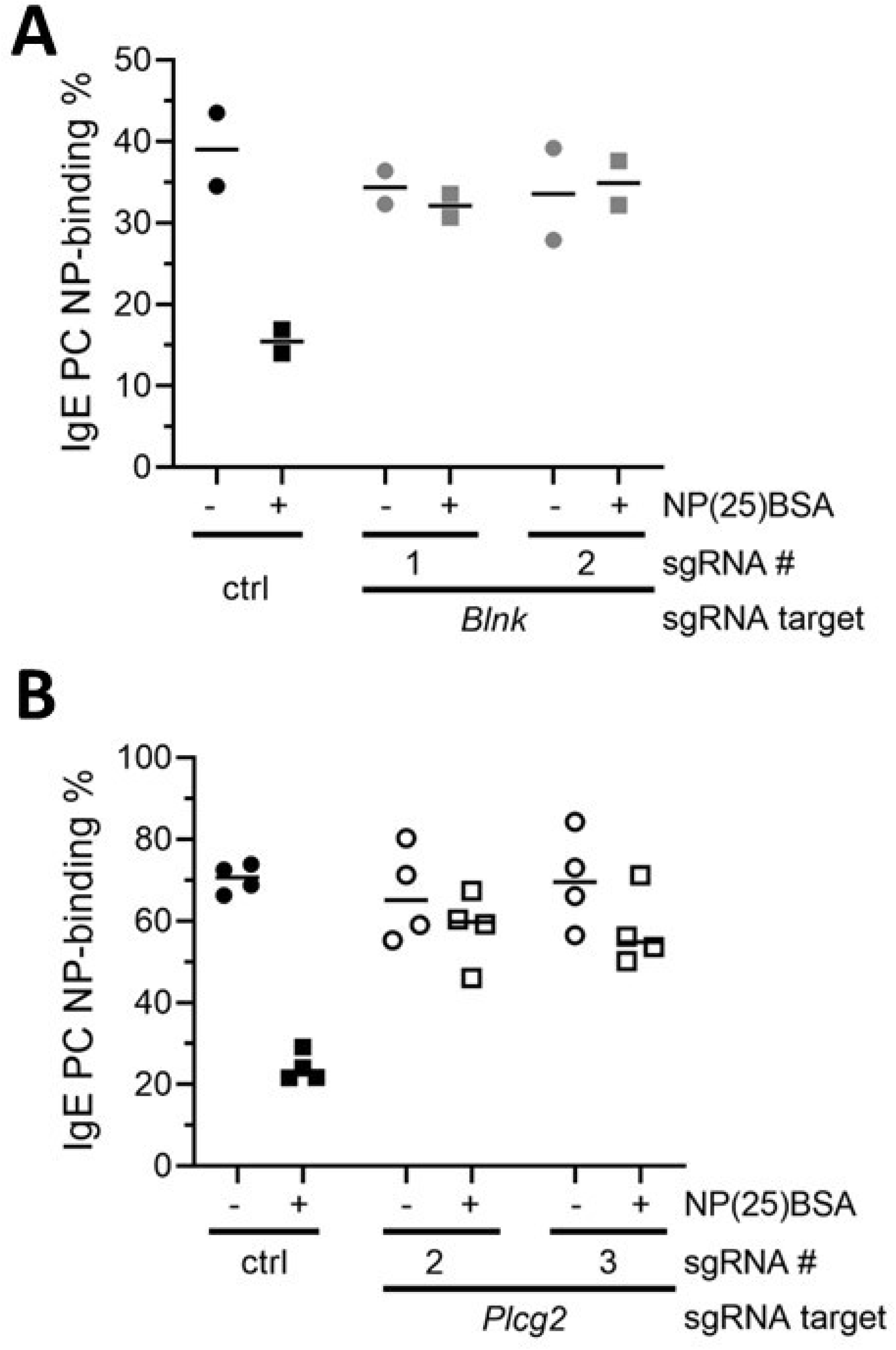
(A-B) Genes were targeted by electroporation with CRISPR-Cas9 ribonucleoproteins as described in Figure 5 and Methods. The proportion of IgE PCs that were NP-binding was determined by flow cytometry. Control conditions either received no guide RNA **(A)** or a non-targeting sgRNA **(B)**. IgE PCs were gated as B220^int^CD138^+^IgE^+^. Dots represent data points derived from cells from individual mice. Bars represent the mean. Results are representative of two independent experiments.

## Notes

### Summary of Updates

1. Figure 1 was revised and split into two figures (Figure 1 and S2) to provide additional quantification graphs and statistics. 2. A new Figure S1 was added to show a representative flow cytometry gating scheme. 3. Figure 6 and the associated supplementary figure were rearranged to new Figures 6 and 7. 4. New data was added showing that cognate antigen exposure in vivo results in a selective reduction in antigen-specific IgE PCs (Figure 7). 5. Modifications were made in the text to incorporate the changes above and clarify the relationship of the work to prior studies. 6. References and mouse nomenclature were updated.

